# Cas1 mediates the interference stage in a phage-encoded CRISPR-Cas system

**DOI:** 10.1101/2024.03.09.584257

**Authors:** Laixing Zhang, Hao Wang, Jianwei Zeng, Xueli Cao, Zhengyu Gao, Zihe Liu, Feixue Li, Jiawei Wang, Yi Zhang, Maojun Yang, Yue Feng

## Abstract

CRISPR-Cas systems are prokaryotic adaptive immune systems against invading phages and other mobile genetic elements, which function in three stages: adaptation, expression and interference. Interestingly, phages were also found to encode CRISPR-Cas systems to antagonize their hosts and establish infection. The *Vibrio cholerae*-infecting ICP1 phage was the first discovered phage with a functional CRISPR-Cas system against an antiphage region in the host genome. Nevertheless, this system lacks a domain essential for recruitment of helicase-nuclease Cas2/3 during target DNA cleavage, and how this system accomplishes the interference stage remains unknown. Here, surprisingly, we found that Cas1, a highly conserved component known to exclusively work in the adaptation stage, also mediates the interference stage through connecting Cas2/3 to the DNA bound-Cascade (Csy) complex of the ICP1 CRISPR-Cas system. A series of structures of Csy, Csy-dsDNA, Cas1-Cas2/3 and Csy-dsDNA-Cas1-Cas2/3 complexes collectively reveal the whole process of Cas1-mediated target DNA cleavage by the ICP1 CRISPR-Cas system. Together, these data support an unprecedented model in which Cas1 mediates the interference stage in a phage-encoded CRISPR-Cas system and also shed light on a unique model of primed adaptation.

## Introduction

Prokaryotic cells have evolved numerous sophisticated defense mechanisms to counteract invading phages and plasmids, including the CRISPR-Cas adaptive immune system ^1–3^, which relies on CRISPR loci and a diverse cassette of CRISPR-associated (Cas) genes. Found in ∼40 % of bacteria and ∼85 % of archaea, CRISPR-Cas systems comprise two broad classes, which are divided into six types (I-VI) and 33 subtypes ^4^. Class 1 systems (types I, III and IV) mainly employ multi-subunit complexes guided by CRISPR RNA (crRNA), and class 2 systems (types II, V and VI) depend on single-subunit Cas proteins ^5^. In general, CRISPR-Cas immunity operates in three stages: adaptation, expression and interference ^6^. During adaptation, fragments of foreign genetic elements are incorporated into the CRISPR array in the host genome as “spacers”, in which process Cas1, Cas2 (or Cas2/3 in type I-F subtype) and in some cases Cas4 ^7^ or certain exonucleases ^8^ are involved. Secondly, the CRISPR array is transcribed into a long transcript (pre-crRNA) which is further processed into crRNAs (the expression stage). In the interference stage, crRNA-guided Cas nucleases, composed of target binding module and target cleavage module, detect invading nucleic acids with a complementary sequence to the spacer (protospacer). These ribonucleoprotein (RNP) complexes first recognize the protospacer adjacent motif (PAM) within the invading nucleic acids, followed by base pairing between the target strand and the spacer, and then the nuclease activity of the RNP complex triggers cleavage of the foreign nucleic acids.

In turn, various tactics are also used by phages to overcome bacterial defense systems ^9–11^. Out of these strategies, surprisingly, phages were also identified to carry functional CRISPR-Cas systems with spacers targeting genomes of their hosts or other phages ^12–14^ and a number of other potential CRISPR-Cas systems ^15–18^. The most intriguing case of this is the utilization of CRISPR-Cas system by the *Vibrio cholerae* serogroup O1-specific ICP1 (the International Centre for Diarrhoeal Disease Research, Bangladesh cholera phage 1)-related phages ^12^ and five other distinct but related *V. cholerae* phages (JSF5, 6, 13, 14, 17 phages) ^18^. *V. cholera* serogroup O1 is the primary cause of cholera, the burden of which could be alleviated by factors including phages in some endemic regions ^12, 19^. Some *V. cholerae* strains rely on a phage inducible chromosomal islands (PICIs) like element (PLE) to protect themselves against the infection by its phages ^20, 21^. Interestingly, out of 26 ICP1-related phages isolated from stool samples of cholera patients, 18 were found to contain a CRISPR-Cas system with spacers against the PLE of the infected *V. cholera* strains ^22^. Moreover, the CRISPR-Cas system from ICP1_2011_A phage has been proved as a fully active one with several hallmarks, such as transcription and processing of the crRNAs, degrading of PLE DNA based on its sequence identity with the CRISPR spacers, and acquisition of new CRISPR spacers targeting the PLE by the derivative phages ^12^.

Typically, both the CRISPR-Cas systems carried by ICP1 and JSF phages belong to the type I-F Csy (CRISPR system yersinia) system ^12, 18^, which recognizes target DNA sequences through a multi-subunit surveillance Cascade (CRISPR-associated complex for antiviral defense) called Csy complex. In contrast to the canonical G-G/C-C PAM used by the type I-F CRISPR-Cas systems in bacteria ^23^, the ICP1/JSF CRISPR-Cas system uses a G-A/C-T PAM adjacent to the protospacers ^12, 18^. Recently, cryo-EM structures of the type I-F Csy complexes from *Pseudomonas aeruginosa* have been reported with and without bound target DNA ^24–26^, shedding light on the architecture of the Csy complex and mechanism of the interference stage, that is, DNA targeting by the surveillance complex. The type I-F CRISPR-Cas system from *P. aeruginosa* (short for *Pae* system hereafter) encodes a ∼350 kDa Csy complex composed of four types of protein and nine subunits (one Cas5f, one Cas8f, one Cas6f and six Cas7f subunits) and crRNA. Following PAM recognition, the target dsDNA (double-stranded DNA) is unwound, rendering hybridization of the target DNA strand with the crRNA spacer and formation of the R-loop by displacing the non-target DNA strand. Importantly, binding of the target DNA induces a ∼180-degree rotation of the C-terminal helical bundle (HB) of the ‘‘large’’ Cas8f subunit of the Csy complex to expose a ‘‘nuclease recruitment helix’’ in Cas8f-HB, which is responsible for further recruitment of the target DNA degrading endonuclease Cas2/3 ^24^. Cas8f-HB is crucial for the type I-F CRISPR-Cas immunity in *P. aeruginosa*, and phages also encode several anti-CRISPR proteins that antagonize the CRISPR-Cas immunity through targeting or mimicking Cas8f-HB, including AcrIF3 ^24^, AcrIF4 ^27^ and AcrIF5 ^28^. In addition, target DNA binding-induced conformational changes of Cascade for recruitment of Cas nucleases have also been demonstrated in many other type I CRISPR-Cas systems ^29–31^.

Sequence analysis reveals smaller ICP1 Cas proteins compared to their *Pae* counterparts, and most interestingly, the ICP1Cas8f protein lacks the HB domain (Figure 1a), representing the most notable difference between the two systems. Therefore, it is unknown how the ICP1 Csy complex recruits the Cas2/3 nuclease for DNA cleavage after recognition of target DNA. Surprisingly, our study uncovers that Cas1, typically involved in adaptation ^32–35^, mediates the interference stage in the ICP1 system. In contrast to its repressive role in *Pae* type I-F system ^35^, Cas1 in ICP1 connects Cas2/3 to target DNA-bound Csy complex, demonstrated through structural, biochemical, and *in vivo* analyses Totally 10 structures of the ICP1 Csy-dsDNA-Cas1-Cas2/3 complex, Csy-dsDNA, Cas1-Cas2/3 and apo Csy showed that target DNA binding induced elongation of the ICP1 Csy backbone, creating space for Cas1-Cas2/3 and exposing the R-loop to Cas2/3’s nuclease active site. Biochemical, *in vivo*, and phage analyses confirm Cas1’s essential role in targeted DNA cleavage. This study pioneers Cas1’s involvement in the interference stage, stimulating the activity of Cas nucleases. Additionally, these structures represent the first of a complete class 1 CRISPR-Cas system from phage origins, providing insights into the evolutionary relationship between bacterial and phage-origin CRISPR-Cas systems.

**Fig. 1.**
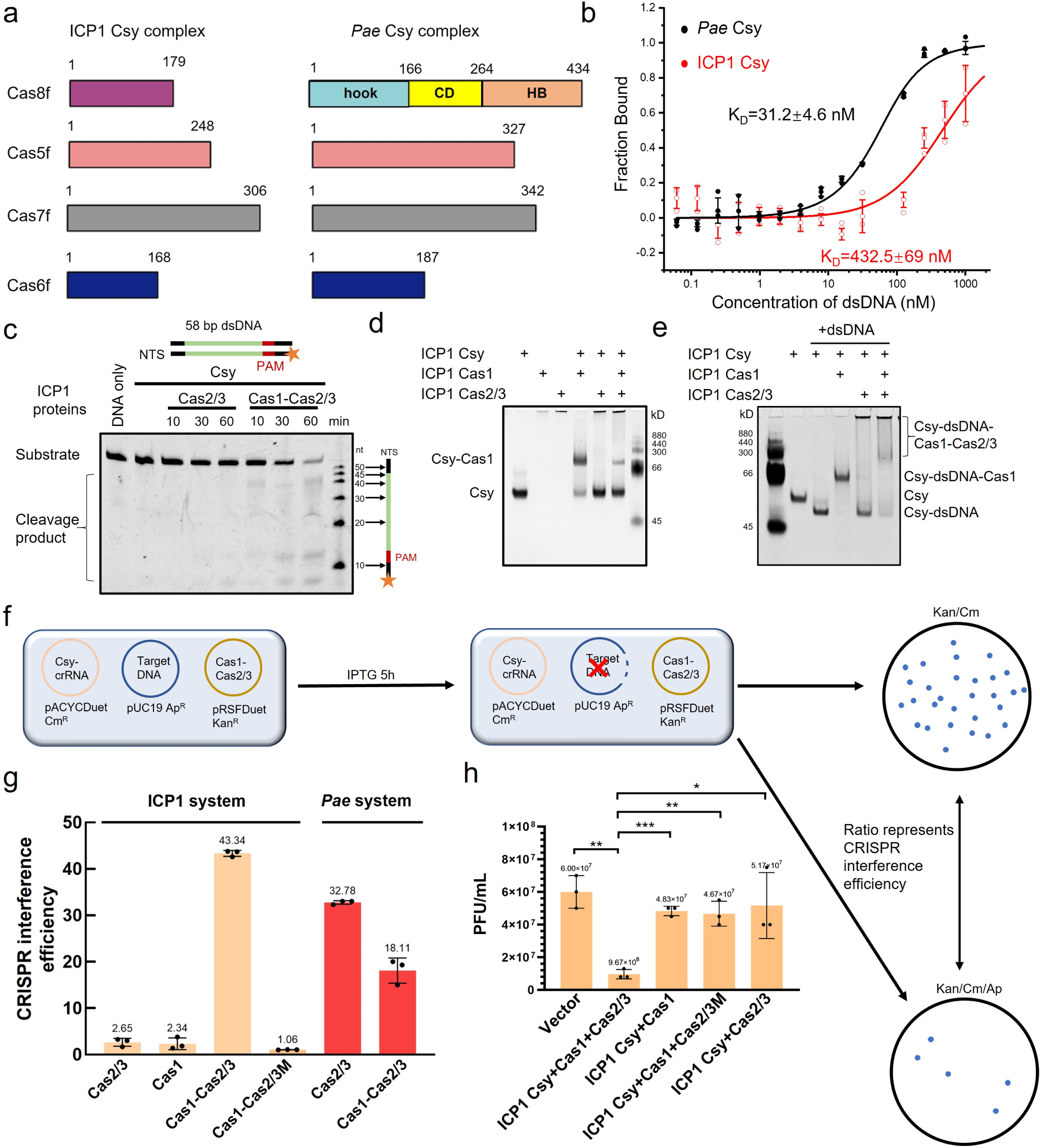
Cas1 mediates target DNA cleavage through connecting Csy-dsDNA complex to Cas2/3. **a,** Comparison of cascade components between the type I-F CRISPR-Cas systems of ICP1 phage and *P. aeruginosa.* Domain architecture of *Pae*Cas8f is marked. **b,** Microscale Thermophoresis (MST) assays to test the binding of target dsDNA to the Csy complex of ICP1 and *Pae*. The binding curves are displayed and fitted to calculate the K_D_ values. The error bars represent the s.d. of each data point calculated from three independent measurements. **c,** *In vitro* DNA cleavage. 0.05 µM dsDNA was preincubated with 0.8 µM Csy complex. Next, Cas2/3 (0.1 µM) or Cas1-2/3 (0.05 µM) with 1 mM ATP and 4 mM MnCl_2_ were added into the reaction system. The reaction was terminated at 10, 30 or 60 min. The products were separated by Urea-PAGE and visualized by fluorescence imaging. **d-e,**. Protein binding assays. 1 µM Csy (**d**) or Csy-dsDNA (**e**) was incubated with 4 µM Cas2/3 only, or first incubated with 2 µM Cas1 followed by adding 4 µM Cas2/3. The mixture was separated by native PAGE and visualized by Coomassie blue R250 staining. **f,** Schematic of the plasmid interference assay. Interference efficiency is measured as the number of colonies on the Kan/Cm plate divided by that on the Kan/Cm/Ap plate. **g,** Plasmid interference of ICP1 and *Pae* CRISPR-Cas system. Mean and s.d. from three independent experiments are shown. Cas2/3M in this figure represents Cas2/3 D112A/D280A. **h,** Plaque assays of the ICP1 CRISPR-Cas system. The experiment has been repeated independently for 3 times.*: p⩽0.033, **: p⩽0.002, ***: p<0.001. PFU: Plaque forming Unit.

## Results

### Cas1 mediates target DNA cleavage in the ICP1 CRISPR-Cas system

The ICP1Cas8f lacks the HB domain and also shows low protein sequence identity with the *Pae*Cas8f (∼12% identity, Figures 1a; Extended Data Figure 1a). Importantly, in the *Pae* system the HB domain is not only essential for Cas2/3 recruitment, but also involved in target DNA binding ^24^. Consistently, ICP1 Csy displays a weaker binding to target DNA compared to *Pae* Csy when they use the same spacer sequence (Figure 1b). However, while ICP1 Csy was able to bind target DNA, as expected, further including Cas2/3 did not result in efficient cleavage of the DNA (Figure 1c), possibly due to lack of the HB domain in ICP1Cas8f. During our research, we accidentally found that the ICP1 Csy complex is capable of binding to Cas1 (Figure 1d), which was not observed in the case of the *Pae* type I-F system (Extended Data Figure 1b). To our knowledge, this binding between Cas1 and a CRISPR Cascade has not been reported in any other CRISPR-Cas types either. Therefore, we moved on to test whether Cas1 has the potential to mediate the binding between Cas2/3 and DNA-bound Csy complex. Interestingly, native PAGE assay showed that while Cas1 is able to bind both Csy and Csy-dsDNA, Cas2/3 can only bind to the Csy-dsDNA-Cas1 complex but not the Csy-Cas1 complex (Figure 1d, e). Notably, binding with dsDNA will increase the overall negative charge in the Csy-dsDNA complex, resulting in a faster migration than Csy alone in the gel (Figure 1e). Therefore, we hypothesized that Cas1 might facilitate the cleavage of target DNA in the ICP1 system. Consistent with our hypothesis, *in vitro* cleavage assay clearly showed that cleavage of target DNA could be greatly enhanced in the presence of Cas1 (Figure 1c). This was also supported by an *in vivo* CRISPR interference assay in *Escherichia coli* we set up (Figure 1f), which showed that only in the presence of Cas1 could the target DNA be efficiently cleaved by Cas2/3 and Csy in the ICP1 system (Figure 1g). In contrast, the *Pae* type I-F CRISPR-Cas system was fully active in target DNA cleavage without Cas1 in the same assay (Extended Data Figure 1c), and including Cas1 even showed some inhibition effects on the activity of the *Pae* system, which is consistent with the previous notion ^35^. As previously reported, Cas1 was not needed for DNA cleavage by the *Pae* CRISPR-Cas system in the *in vitro* assay either (Extended Data Figure 1c). Finally, we incorporated the ICP1 type I-F system with a synthetic CRISPR array containing a spacer directed to the transcribed strand of the mrh24 and uvsX gene transcripts of the dsDNA phage T4 into *E. coli* BL21. Consistently, end point assays showed that inclusion of Cas1 greatly enhances the anti-phage activity of Csy and Cas2/3 in the ICP1 system (Figure 1h and Extended Data Figure 2). Taken together, these results demonstrate that in the ICP1 CRISPR-Cas system, Cas1 unexpectedly mediates target DNA cleavage through connecting Csy-dsDNA complex to Cas2/3.

### Overall structure of the ICP1 Csy complex

To elucidate ICP1Cas1’s role in recruiting Cas2/3 to the Csy-dsDNA complex, we purified the Csy-dsDNA-Cas1-Cas2/3 complex to homogeneity through affinity and size-exclusion chromatography (Extended Data Figure 1d,e). Structures of ICP1 Csy, Csy-dsDNA, Cas1-Cas2/3, and Csy-dsDNA-Cas1-Cas2/3 were determined via X-ray crystallography and single-particle cryo-EM (Supplementary Table 1, 2; Extended Data Figure 3, 4). Since both ICP1 Csy and Csy-dsDNA show structural features different from those of the *Pae* complexes, we will first introduce the structure of ICP1 Csy and its conformational change upon DNA binding. We solved the structure of ICP1 Csy through X-ray crystallography at 3.0 Å resolution, which is the first crystal structure of a Cascade of type I-F CRISPR-Cas systems. During the study, we also solved the crystal structures of ICP1 Cas8f/5f subcomplex and Cas7f, respectively (Supplementary Table 1). Similar to the *Pae* Csy complex, the ICP1 Csy is also composed of nine Cas proteins and a single 60-nt crRNA with 28-nucleotide repeat sequence and 32-nucleotide spacer sequence (Figure 2a). Due to lack of the Cas8f-HB, the ICP1 Csy is not a “closed” ring-like structure as the *Pae* Csy. In all the independent structural studies into the *Pae* Csy complex up to now, the densities for Cas6f and the 3’ hairpin of the crRNA are not well resolved. However, although weaker than the other regions, the density for this region still allows us to *de novo* build the model of ICP1Cas6f and the 3’ hairpin of the crRNA in our crystal structure (Extended Data Figure 5a). This also indicates that flexibility of the linkage tethering Cas6f to the rest of the complex might be diverse among Csy complexes from different species. Consistent with *Pae* Csy complex and the other class 1 complexes, the ICP1 Csy also exhibits crRNA distortions (kinks) along the Cas7f backbone at regular 6-nucleotide intervals (Extended Data Figure 5b). The torsion angles at each kink are similar between ICP1 Csy and *Pae* Csy complexes, resulting in their similar helical pitches, which are substantially tighter than those of the type I-E and III-B complexes. Specifically, the ICP1 Csy backbone adopts a slightly tighter pitch (∼75 Å) than the *Pae* Csy backbone (∼80 Å).

**Fig. 2.**
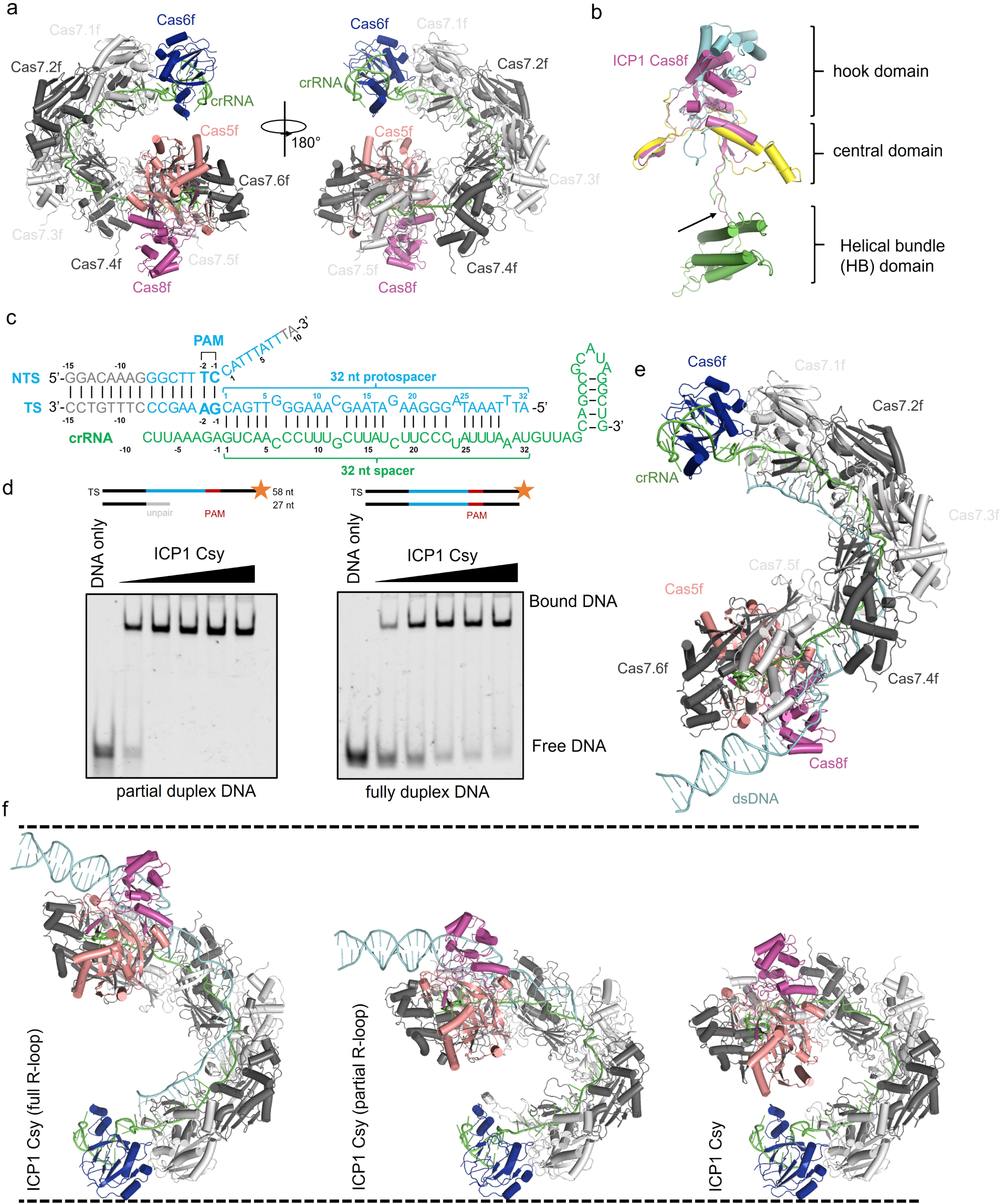
Structure of the ICP1 Csy complex and Csy-dsDNA complex. **a,** Overall crystal structure of the ICP1 Csy complex. The ICP1 Csy complex also displays a seahorse shape, with the typical head (Cas6f), backbone (Cas7f), and tail (Cas5f and Cas8f) subunits of class 1 CRISPR-Cas complexes. **b,** Structural superimposition of ICP1Cas8f and *Pae*Cas8f (PDB code: 6B45). **c,** Schematic drawing of the partial R-loop construct with dsDNA and crRNA in the Csy-dsDNA structure. Nucleotides with no clear density are shown in gray. **d,** EMSA used to test the binding affinities of the ICP1 Csy complex for partial duplex dsDNA and full duplex dsDNA with FAM labeled on the 3′ end of TS DNA. Reactions were performed with 0.1 μM dsDNA and the ICP1 Csy complex concentrations of 0.1, 0.2, 0.4, 0.8 and 1.6 μM following the order indicated by the black triangle. **e,** Cryo-EM structure of ICP1 Csy-dsDNA (partial duplex) complex (colored as in **a**). Partial duplex dsDNA is colored in cyan. **f,** Comparison of the structures of ICP1 Csy, ICP1 Csy-dsDNA (partial R-loop) and ICP1 Csy–dsDNA (full R-loop), with the two dashed lines as a guide to highlight the elongation of the complex.

The stable Cas5f-Cas8f complex constitutes the tail of the ICP1 Csy complex with the S-shaped 5’ handle region of the crRNA. Compared to *Pae*Cas8f which consists of an N-terminal “hook” domain, a central domain (CD) and a C-terminal HB domain, ICP1Cas8f is more compact with regions corresponding to the N-terminal “hook” and central domain of *Pae*Cas8f, although both domains show structural variations between ICP1Cas8f and *Pae*Cas8f (Figure 2b). As aforementioned, the most striking difference between ICP1Cas8f and *Pae*Cas8f is that ICP1Cas8f lacks the HB domain, but only has a loop extruding out from the main domain at the C-terminus (Figure 2b). Although the sequence identity between ICP1Cas5f and *Pae*Cas5f is low (∼17%) and ICP1Cas5f is almost 80 residues shorter than *Pae*Cas5f in sequence, the overall fold of the two proteins is conserved (Extended Data Figure 5c). In the Cas7f backbone, consistent with *Pae*Cas7f and other Cas7 structures, the ICP1Cas7f also has the fingers-, palm-, web-, and thumb-shaped domains, with a “right-hand” morphology (Extended Data Figure 5d). The Cas7f fold is conserved between ICP1Cas7f and *Pae*Cas7f with an average RMSD (root-mean-square deviation) of ∼1.0 Å. They mainly differ in their thumb and web domains, which are shorter in ICP1Cas7f and to be described below (Extended Data Figure 5d). The Cas6f fold is also conserved between ICP1 Csy and *Pae* Csy (Extended Data Figure 5e). Taken together, ICP1 Csy exhibits typical features of a class 1 CRISPR-Cas complex, but also displays its own distinct characteristics.

### Structural overview of the DNA-bound ICP1 Csy complex

Then we moved on to investigate the conformational change of ICP1 Csy upon target DNA binding. Since ICP1 Csy displays a weaker binding to target DNA than *Pae* Csy (Figure 1b), we designed a target DNA with 17 base pairs of PAM-proximal DNA duplex, a 32-nt crRNA-complementary protospacer in the target DNA strand (TS), and a 10-nt overhang at the 3’ end of the non-target DNA strand (NTS) (Figure 2c), to avoid the reannealing of the protospacer region of target DNA and thereby get a better binding by the ICP1 Csy. As expected, EMSA (electrophoretic mobile shift assay) showed that ICP1 Csy displays a higher binding affinity to this partial duplex DNA than the canonical dsDNA (Figure 2d). Then we reconstituted the ICP1 Csy-dsDNA complex with this partial duplex DNA and solved its cryo-EM structure (Figure 2e; Extended Data Figure 5). In the structure, almost complete sequence of both TS and NTS of the target DNA could be modeled. As was observed in the *Pae* Csy-dsDNA complex ^24, 25^, the DNA-bound ICP1 Csy complex also displays a significant conformational change compared to the apo Csy, the most notable feature of which is the elongation of the ICP1 Csy backbone by ∼32.5 Å caused by rigid-body rearrangement (Figure 2f). Interestingly, during the cryo-EM studies into the ICP1 Csy-dsDNA-Cas1-Cas2/3 complexes (to be described below), particles corresponding to Csy-dsDNA complex (∼20% of the total particles), which represented partially assembled complex were also observed. These particles were classified into two classes and their structures were solved respectively (Supplementary Table 2; Extended Data Figure 3, 4). Notably, while the dsDNA used in reconstituting this whole complex was complete duplex DNA, one out of the two structures was very similar with the Csy-dsDNA structure using the partial duplex DNA (Extended Data Figure 6a). In this Csy-dsDNA structure, the complete 32-nt protospacer region of TS could also be modelled, representing a state with a full R-loop. In the other solved Csy-dsDNA structure using complete duplex DNA, interestingly, only 16 nucleotides in the protospacer region of TS could be modelled, suggesting that it represents an intermediate state before exhibiting a full R-loop. Correspondingly, the overall length of this structure was between those of the apo ICP1 Csy and Csy-dsDNA structure with 32-nt modelled protospacer region (Figure 2f, Supplementary Video 1).

In the ICP1 Csy-dsDNA structure, the duplex region of the target DNA is also sandwiched between the Cas8f, the thumb region of Cas5f and Cas7.6f subunit (Figure 2e). The target and non-target DNA strands separate immediately at the protospacer region next to the PAM site (i.e., G(-1)A(-2)^TS^ and C(-1)T(-2)^NTS^, Figure 2e; Extended Data Figure 6b). Similar to the *Pae* Csy-dsDNA structure, a loop in the Cas8f subunit (residues 141-148, sequence TQKLGGSN) is wedged into the dsDNA, and sterically block a 1^TS^-1^NTS^ base pair, with Cas8f-N148 stacking with PAM G(-1)^TS^ (Extended Data Figure 6b). Besides, Cas8f-S147 interacts with the phosphate group of G(-1)^TS^ with a hydrogen bond and Cas8f-Q150 interacts with the base of G (-1)^TS^ via a hydrogen bond. Cas5f-I88 also hydrogen bonds with the phosphate group of G(-1)^TS^ (Extended Data Figure 6b). These observations are consistent with the *in vivo* results, which indicated that the G(−1) of the PAM sequence is critical for target DNA recognition ^22^. Next to the PAM, the two DNA strands are separated immediately, where the TS forms a discontinuous heteroduplex with the crRNA spacer, with one kinked-off nucleotide at every 6^th^ position (Extended Data Figure 6c). After PAM, the NTS is attached to a positively charged groove formed mainly by the Cas8f subunit (Extended Data Figure 6d), which was observed in all the three ICP1 Csy-dsDNA structures. This is also consistent with the cases in other type I effector complexes, where the NTS is mainly flexibly attached to the effector Cascade. The ICP1 Csy-dsDNA structure also helps to explain the weaker DNA binding affinity of ICP1 Csy as compared to *Pae* Csy. Previous studies have revealed two major roles of the Cas8f HB domain of *Pae* Csy in facilitating target DNA binding ^24^. On the one hand, after ∼180° rotation, the HB domain folds over the TS and further binds the ‘thumbs’ of Cas7.2f and Cas7.3f from one side, and the ‘webs’ of them from the opposite site of the TS, thus completely encircling the target DNA strand (Extended Data Figure 6e). On the other hand, the rotated HB together with Cas5f subunit forms a R-loop binding channel (RBC) to accommodate the NTS ^24^. Taken together, Cas8f-HB prevents the reannealing of target dsDNA through these two approaches. However, the ICP1 Cas8 lacks the HB domain and the ICP1 Cas7 has shorter “thumbs” and “webs” (Extended Data Figure 5d). Specifically, ICP1Cas7f has fewer positively charged residues than *Pae*Cas7f within the “thumb” and “web” regions. K58 and K60, two residues revealed to be critical for dsDNA binding by the *Pae* Csy ^26^, are also lacking in the ICP1Cas7f (Extended Data Figure 6f). Therefore, lacking Cas8f-HB and the distinct structures in the Cas7f “thumb” and “web” may collectively result in the lower binding efficiency to target DNA by the ICP1 Csy. Consistently, a very recent study also showed that the weak binding to ICP1 Csy by target dsDNA is caused by its tendency to reanneal after unwinding following PAM recognition ^36^.

### Architecture of the ICP1 Csy-dsDNA-Cas1-Cas2/3 complex

The most striking feature of the ICP1 type I-F system lies in that Cas1 is essential for the interference stage. The architecture of Csy-dsDNA-Cas1-Cas2/3 complex was captured in two different states in the cryo-EM study, which represent a fully assembled complex and approximately half of it, respectively (Supplementary Table 2; Extended Data Figure 4). Here we will first describe the architecture of the fully assembled Csy-dsDNA-Cas1-Cas2/3 complex (Figure 3a, b, Supplementary Video 2). In this huge complex, a Cas1-Cas2/3 complex with a stretch of dsDNA bound is in the center containing one Cas2/3 dimer and two Cas1 dimers, the ratio of which is consistent with the low resolution models of Cas1-Cas2/3 complex of bacterial type I-F CRISPR-Cas system ^35, 37^ and structures of Cas1-Cas2 complex of other systems ^34, 38, 39^. Two Csy-dsDNA complexes are flanking the central Cas1-Cas2/3 complex (Figure 3a, b), so that the hetero-24-mer complex exhibited 2-fold rotational symmetry, with a total molecular weight up to 1 MDa. The structure can be divided into two Csy-dsDNA-Cas1-Cas2/3 subcomplexes, each forming a closed ring with one Csy-dsDNA, one Cas2/3 and a Cas1 dimer (Figure 3a, b). The aforementioned “half” form of the complex contains one such subcomplex plus the other Cas1 dimer and Cas2/3 protein, the density of whose Cas3 domain was not visible (Figure 3c), suggesting flexibility of this domain. In the complex structure, the head (Cas6f) and tail (Cas5f) region of Csy-dsDNA interacts with the exterior Cas1 protomer (Cas1.2) and Cas2/3, respectively (Figure 3a, b), so that the separated NTS is extended into the Cas2/3 nuclease. Importantly, structures of the Csy-dsDNA-Cas1-Cas2/3 complex also explain why Cas2/3 can only bind to the Csy-dsDNA-Cas1 complex but not the Csy-Cas1 complex (Figure 1d, e): Superimposition of the apo Csy onto the Csy within the complex structure at the Cas6f subunit showed severe steric clash between Cas2/3 and the Csy backbone, especially the tail (Cas5f/8f) region (Extended Data Figure 7a), indicating that only by elongating the Csy complex through target DNA binding could create enough space to accommodate the Cas1-Cas2/3 complex. On the other hand, structural comparison between the Csy-dsDNA complex and that within the Csy-dsDNA-Cas1-Cas2/3 complex reveals only slight conformational changes mainly at the Cas6f part (Extended Data Figure 7b).

**Fig. 3.**
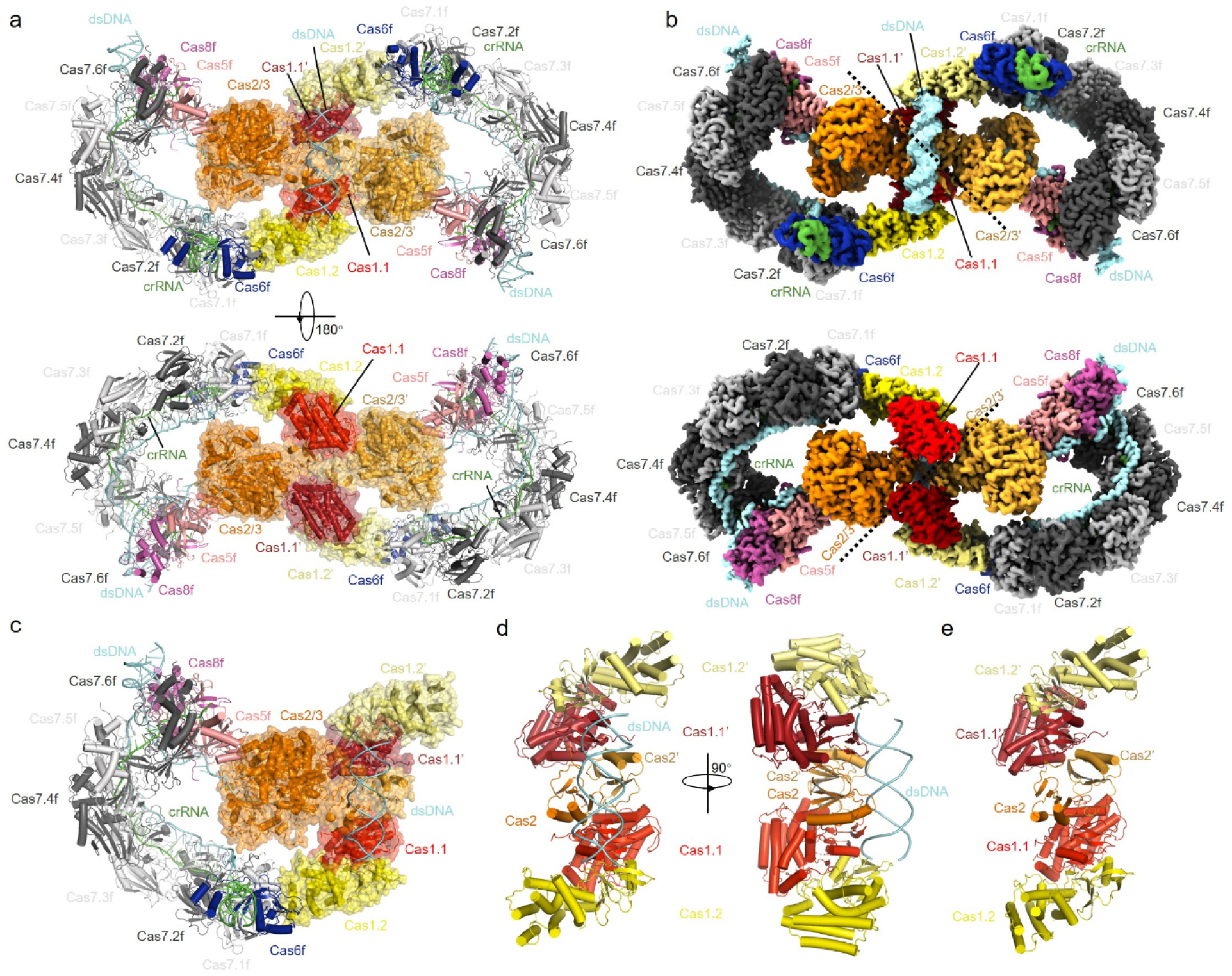
Cryo-EM structure of the ICP1 Csy-dsDNA-Cas1-Cas2/3 complex. **a,** Overall structure of the ICP1 Csy-dsDNA-Cas1-Cas2/3 complex in cartoon representation with each subunit colored as in Figure 2e. Two views are shown. **b,** Cryo-EM map of the ICP1 Csy-dsDNA-Cas1-Cas2/3 complex with each subunit color-coded. Two views same as in **a** are shown. **c,** “Half” form of the ICP1 Csy-dsDNA-Cas1-Cas2/3 complex with each subunit color-coded. **d,** Cryo-EM structure of the ICP1 Cas1-Cas2/3-dsDNA complex with no density for the Cas3 domain. Two views are shown. **e,** Cryo-EM structure of the ICP1 Cas1-Cas2/3 complex with no density for the Cas3 domain.

To explore potential conformational changes in the Cas1-Cas2/3 complex upon Csy-dsDNA binding, cryo-EM analysis revealed particles corresponding to Cas1-Cas2/3 complexed with dsDNA. However, no density for the Cas3 domain was observed (Figure 3d; Extended Data Figure 4), suggesting its flexibility. We further purified the Cas1-Cas2/3 complex and solved its structure (Supplementary Table 2; Extended Data Figure 4), yet still observed no density for the Cas3 domain (Figure 3e), while the sample gel indicates the integrity of the Cas2/3 protein (Extended Data Figure 8a), further indicating its high flexibility. Structural superimposition of the Cas2 dimer of the apo and DNA-bound ICP1 Cas1-Cas2 complex shows that the two Cas1 dimers rotate a bit in either clockwise (Cas1.1/2) or anti-clockwise (Cas1.1’/2’) directions upon DNA binding (Extended Data Figure 8b). In detail, Cas1.1/2 rotate 9.60 degrees and move along the rotation axis 1.19 Å. Cas1.1’/2’ rotate 11.46 degrees and move along the rotation axis 1.10 Å. Furthermore, superposition of the Cas2 dimer of the DNA-bound Cas1-Cas2 and that in the Csy-dsDNA-Cas1-Cas2/3 complex reveals that the two Cas1 dimers rotate greatly in either anti-clockwise (Cas1.1/2) or clockwise (Cas1.1’/2’) directions upon binding to Csy-dsDNA (Figure 4e). In detail, Cas1.1/2 rotate 16.74 degrees and move along the rotation axis 0.69 Å. Cas1.1’/2’ rotate 16.41 degrees and move along the rotation axis 0.78 Å. During this process, while Cas2/2’ and Cas1.2/1.2’ undergo mainly rigid-body movements (RMSD of ∼0.53 Å and ∼0.46 Å between the DNA-bound Cas1-Cas2 and that in the Csy-dsDNA-Cas1-Cas2/3 complex, respectively), Cas1.1/1.1’ exhibits significant conformational change (RMSD of ∼1.56 Å), especially in helices α3-5 (Extended Data Figure 8c).

**Fig. 4.**
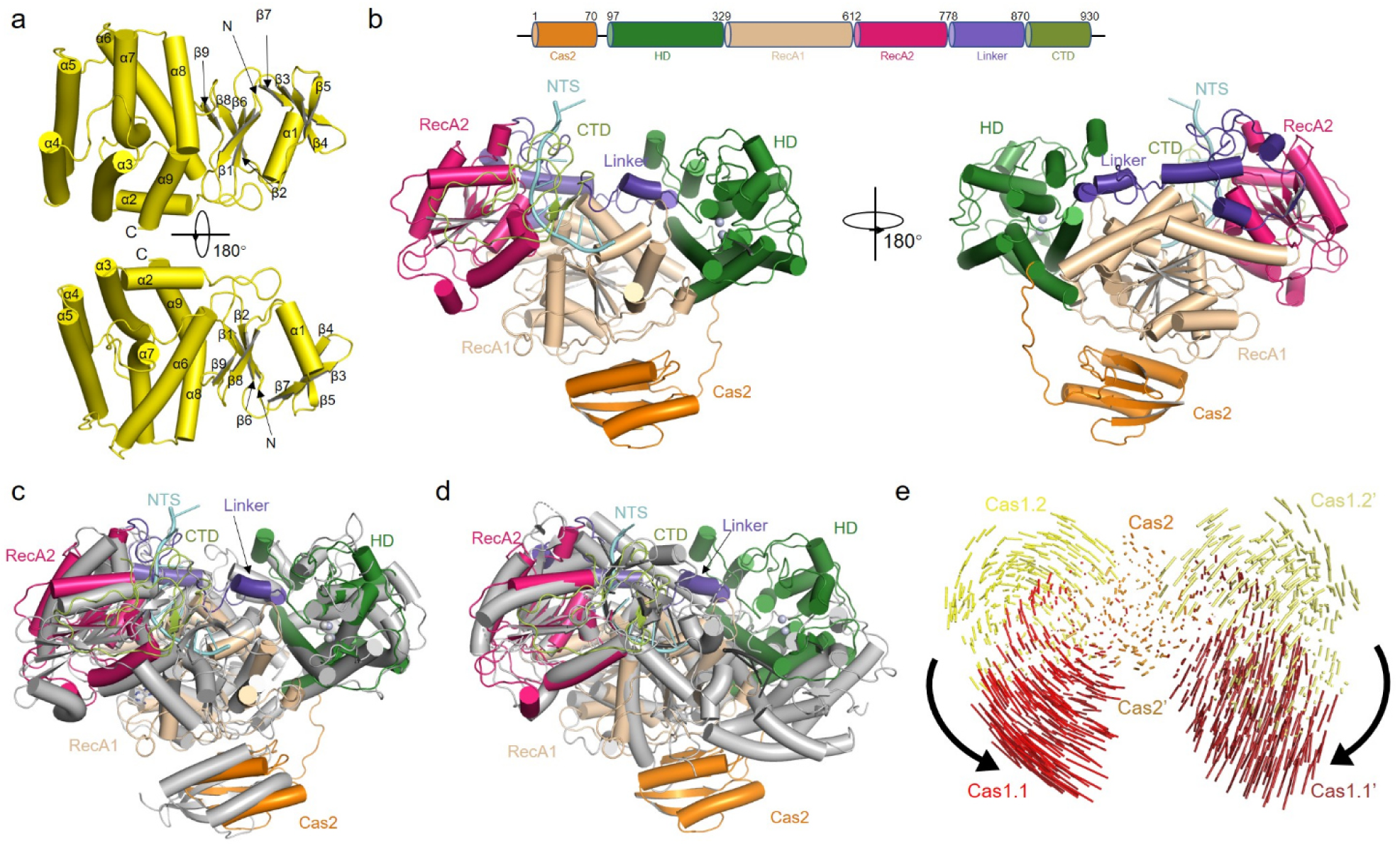
Structural plasticity of the ICP1 Cas1-Cas2/3 complex. **a,** The structure of ICP1Cas1 is shown in cartoon model. Two views are shown and secondary structures are labelled. **b,** Domain organization and structure of ICP1Cas2/3 shown in cartoon. Bound NTS in the Cas2/3 protein is colored in cyan. Cas2 domain (1-78), an active histidine-aspartate (HD) domain (96-329 residues), RecA1 domain (330-612 residues), RecA2 domain (613-778 residues), a linker (779-869 residues), and a C-terminal domain (CTD, 870-930 residues). **c,** Structural alignment of ICP1Cas2/3 and *Pae*Cas2/3 (PDB code: 5B7I, gray). **d,** Structural alignment of ICP1Cas2/3 and *Tfu*Cas3 (PDB code: 4QQX, gray). **e,** Structural comparison between ICP1 Cas1-Cas2 in the dsDNA-bound form and that in the Csy-dsDNA-Cas1-Cas2/3 complex structures. The Cas2 dimer is superimposed. Vector length correlates with the domain motion scale. The black arrows indicate domain movements within the Cas1-Cas2 complex upon Csy-dsDNA binding.

### Structure of the ICP1 Cas1-Cas2/3 complex and its structural flexibility

Since the structures of ICP1Cas2/3 and the Cas1-Cas2/3 complex were both solved for the first time and they both show structural features different from those of their *Pae* counterparts and of other systems, next we will introduce the characteristics of ICP1Cas1, Cas2/3 and Cas1-Cas2/3 complex. Purified ICP1Cas1 also forms a dimer as other Cas1 proteins (Extended Data Figure 8d) and displays canonical Cas1 fold with an N-terminal β-strand domain and a C-terminal α-helical domain (Figure 4a). Comparison of ICP1Cas1 in the Csy-dsDNA-Cas1-Cas2/3 complex with Cas1 in its apo form (PDB code: 4W8K) does not reveal marked conformational changes, with an RMSD of ∼0.87 Å (Extended Data Figure 8e). The most similar structural homolog returned by the Dali server is the *Pae*Cas1 protein (Extended Data Figure 8f) with an RMSD of ∼1.18 Å. The ICP1Cas2/3 also displays conserved features of Cas2 and Cas3 proteins (Figure 4b). *Pae*Cas2/3 is the most similar structure returned by the Dali server (Figure 4c). While the Cas2 domains of ICP1Cas2/3 and *Pae*Cas2/3 are in different orientations in the structural alignment (Figure 4c), their structures themselves are similar with each other (Extended Data Figure 8g, RMSD of ∼0.78 Å). Typically, the active site conformation of the HD nuclease and RecA1 helicase domains are highly conserved between ICP1Cas2/3 and *Pae*Cas2/3 (Extended Data Figure 8h and 8i). Consistently, mutation of D112/D280 abolished the activity of the ICP1 CRISPR-Cas system in both *in vivo* plasmid interference (Figure 1g) and phage assays (Figure 1h and Extended Data Figure 2). However, the conformations of the RecA2 domain and CTD are markedly different between ICP1Cas2/3 and *Pae*Cas2/3 (Extended Data Figure 8j), which can result from the NTS DNA caught in the ICP1Cas2/3 but not the *Pae*Cas2/3 structure. Supporting this, the conformations of the RecA2 and CTD of ICP1Cas2/3 are more similar to those of Cas3 from *Thermobifida fusca* (*Tfu*Cas3, PDB: 4QQX), which also captures DNA (Figure 4d). However, the HD domains of ICP1Cas2/3 and *Tfu*Cas3 are in different orientations (Figure 4d), which might result from the different situations the two proteins are in. That is, the ICP1Cas2/3 is in the Csy-dsDNA-Cas1-Cas2/3 complex and *Tfu*Cas3 is in the free form ^40^.

The structure of Cas1-Cas2/3 complex within the Csy-dsDNA-Cas1-Cas2/3 complex also represents the first atomic structure of Cas1-Cas2/3 of phage CRISPR-Cas systems. Very recently, a high-resolution structure of the type I-F Cas1-2/3 complex from *Pseudomonas aeruginosa* has been reported ^41^. The structural comparison of ICP1 Cas1-2/3 and *P. aeruginosa* Cas1-2/3 shows that they have significant conformational differences due to their different stages (Extended Data Figure 8k).

In both apo and DNA-bound structures of ICP1 Cas1-Cas2/3 complex, Cas3 domain exhibited high flexibility, with no density observed. Unlike bacterial Cas1-Cas2/3 models where Cas1.1 and Cas1.2 interact with Cas2/3 ^35, 37^, in ICP1 Cas1-Cas2/3 complex, Cas3 domain was visible only within the Csy-dsDNA-Cas1-Cas2/3 complex, with only Cas1.1 interacting with Cas2/3 (Figure 3a). This also suggests the different functions of bacterial Cas1-Cas2/3 and ICP1 Cas1-Cas2/3, since the former only participates in adaptation and the latter functions both in adaptation and interference. When the structure of ICP1 dsDNA-Csy-Cas1-2/3 is aligned to the model of *P. aeruginosa* dsDNA-Csy-Cas1-2/3 at their Csy complex (Extended Data Figure 9), ICP1 Cas2/3 and *Pae*Cas2/3 are located at similar position, but the distance between the mass centers of ICP1 Cas2/3’ and *Pae*Cas2/3’ is 98.52 Å. It suggests that the conformation of ICP1 Cas1-Cas2/3 might change dramatically from the adaptation stage to the interference stage. The flexible Cas3 domain might be responsible for this conformation change. Density for dsDNA above Cas1-Cas2 domains was observed (Figure 3b), likely representing non-specifically bound dsDNA used in incubation, distinct from *E. coli* DNA, as apo Cas1-Cas2/3 structures lacked DNA densities in this region (Figure 3e).

### ICP1Cas1 connects Cas2/3 to the Csy-dsDNA complex

In the Csy-dsDNA-Cas1-Cas2/3 complex, the interior Cas1 protomer (Cas1.1/1.1’) binds to Cas2/3 and the exterior Cas1 protomer (Cas1.2/1.2’) mainly binds to the Cas6f subunit of the Csy complex, through which the Cas1 dimer connects Cas2/3 to the Csy-dsDNA complex (Figure 3a). This also explains why ICP1Cas1 is able to bind both Csy and Csy-dsDNA (Figure 1d, e), because structural comparison between Csy and Csy-dsDNA reveals no conformational change of the Cas6f subunit (Extended Data Figure 10a). In ICP1Cas1.2, α5 and the loop linking α5 and α6 (named L56) is involved in binding to the Cas6f subunit, forming hydrogen bond and electrostatic interactions (Figure 5a). Notably, probably due to the interactions from Cas1.2, the region of the Cas6f and the crRNA 3’ hairpin in the structure of Csy-dsDNA-Cas1-Cas2/3 complex shows the clearest density among the Cryo-EM structures of Csy in this study (Extended Data Figure 10b), in contrast to the weak density in this region of *Pae* Csy-dsDNA structures ^24, 25^. In the interface between Cas1.2 and Cas6f, the involved residues of Cas1.2 are primarily at the C-terminus of α5 and L56 (Figure 5b). Importantly, while the E187G mutation of Cas1 does not interfere with the formation of the Cas1-Cas2/3 complex (Extended Data Figure 1f), the mutation severely decreased the activity of the ICP1 system in the *in vivo* plasmid interference (Figure 5c), phage assay (Figure 5d and Extended Data Figure 2) and *in vitro* (Figure 5e) DNA cleavage assay.

**Fig. 5.**
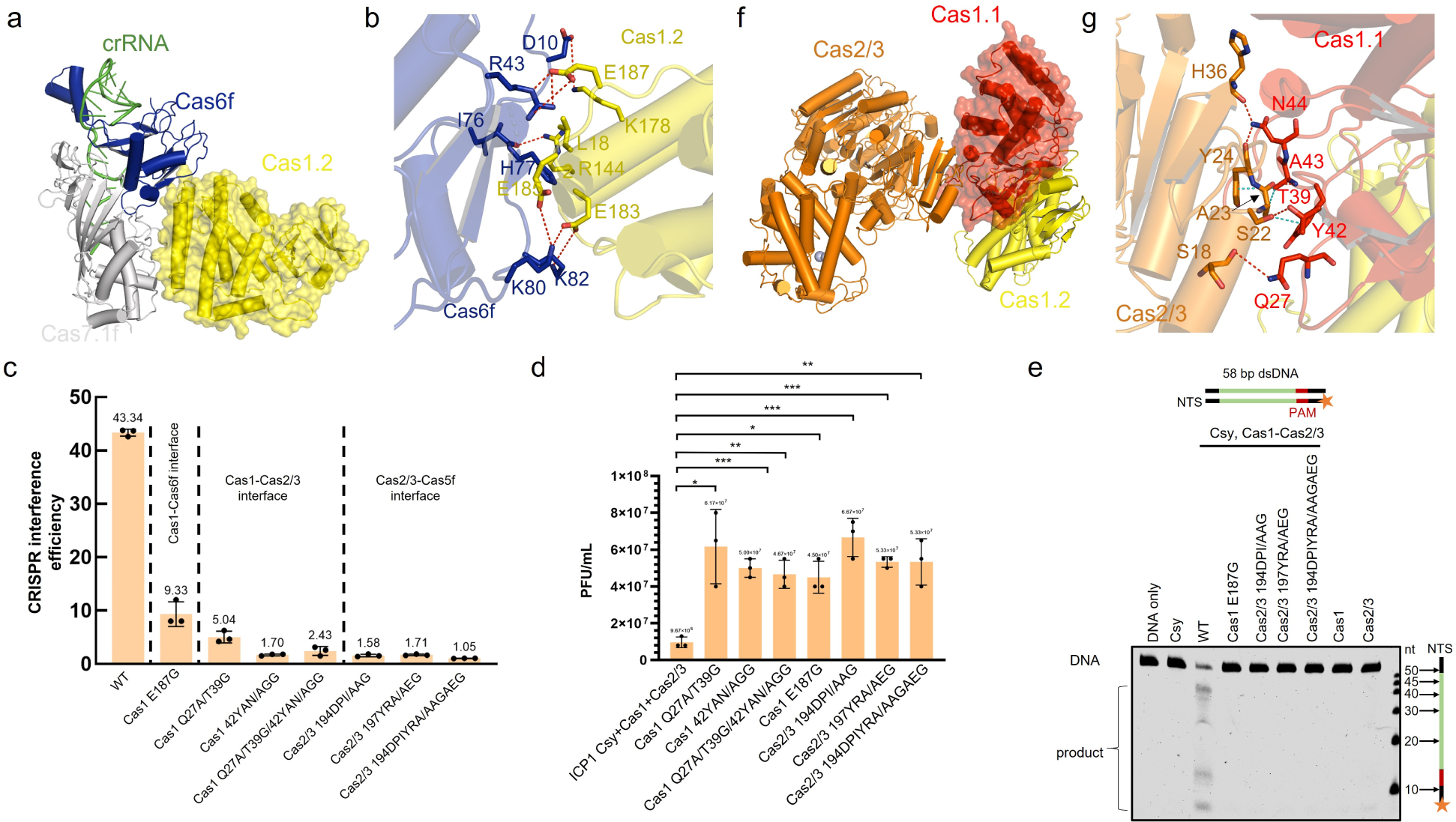
The ICP1Cas1 dimer connects Cas2/3 with the Csy-dsDNA complex. **a,** ICP1Cas1.2 mainly interacts with the Cas6f subunit. The proteins are colored as in Figure 3a. **b,** Detailed interactions between Cas1.2 and Cas6f. Hydrogen bond and electrostatic interactions are shown as red dashed lines. K178/L182/E183/E185/E187 of Cas1.2 interact with D10/R43/H77/K80/K82 of Cas6f. **c,** Plasmid interference of the ICP1 CRISPR-Cas system. Mean and s.d. from three independent experiments are shown. Cas1 42YAN/AGG represents Cas1 Y42A/A43G/N44G. Other mutants with mutations of successive residues are named similarly, in which the first number indicates the number of the first mutated residue. **d,** Plaque assays of the ICP1 CRISPR-Cas system. The experiment has been repeated independently 3 times. *: p⩽0.033, **: p⩽0.002, ***: p<0.001. PFU: Plaque forming Unit. **e,** *In vitro* DNA cleavage assay. 0.05 µM dsDNA was preincubated with 0.8 µM Csy complex. Next, Cas1-Cas2/3 (0.05 µM) or its mutants with 1 mM ATP and 4 mM MnCl_2_ were added into the reaction system. The reaction was terminated at 60 min. The products were separated by Urea-PAGE and visualized by fluorescence imaging. **f,** ICP1Cas1.1 interacts with Cas2/3 and Cas1.2. The proteins are colored as in Figure 3a. **g,** Detailed interactions between Cas1.1 and Cas2/3. Hydrogen bond and electrostatic interactions are shown as red dashed lines, and hydrophobic interactions are represented as teal dashed lines.

On the other hand, the Cas1.1 protomer in the Cas1 dimer mainly interacts with the Cas2/3 protomer in the same subcomplex, apart from binding to Cas1.2 (Figure 5f). Specifically, hydrogen bond and electrostatic interactions are involved, between Q27/T39/N44 of Cas1.1 and S18/S22/H36 of Cas2/3 (Figure 5g). In addition, Y42/A43 of Cas1.1 form hydrophobic and π-alkyl interactions with A23/Y24 of Cas2/3. Detailed interactions are presented in Figure 5g. Consistently, mutations of the interface residues in the Cas1.1-Cas2/3 interface severely hinder the formation of the Cas1-Cas2/3 complex (Extended Data Figure 1f). These mutations also inactivated the ICP1 CRISPR-Cas system in both the *in vivo* (Figure 5c) and phage assays (Figure 5d and Extended Data Figure 2). Interestingly, hydrogen bond and electrostatic interactions are also found between Cas1.1 and the Cas3 domain of Cas2/3’ in the other subcomplex (Extended Data Figure 10c, d). However, these interactions might only exist and be stable in the context of the fully assembled Csy-dsDNA-Cas1-Cas2/3 complex, because there is no density for the Cas3 domain of Cas2/3’ in the “half” form of the Csy-dsDNA-Cas1-Cas2/3 complex (Figure 3c). This also supports that the presence of Csy-dsDNA in the complex can stabilize the Cas3 domain of Cas2/3.

### ICP1Cas2/3 binds to Cas5f and captures NTS

Notably, in contrast to *Pae*Cas2/3 which is recruited to the Csy complex through the HB domain of Cas8f, ICP1Cas2/3 binds to the Cas5f subunit (Figure 3a and 6a,b,c) with a buried surface of 833.5 Å^2^. Consistently, mutations of the Cas2/3 residues which interact with Cas5f, markedly decreased the activity of the ICP1 system in the *in vivo* plasmid interference (Figure 5c), phage assay (Figure 5d and Extended Data Figure 2) and *in vitro* DNA cleavage assay (Figure 5e).

**Fig. 6.**
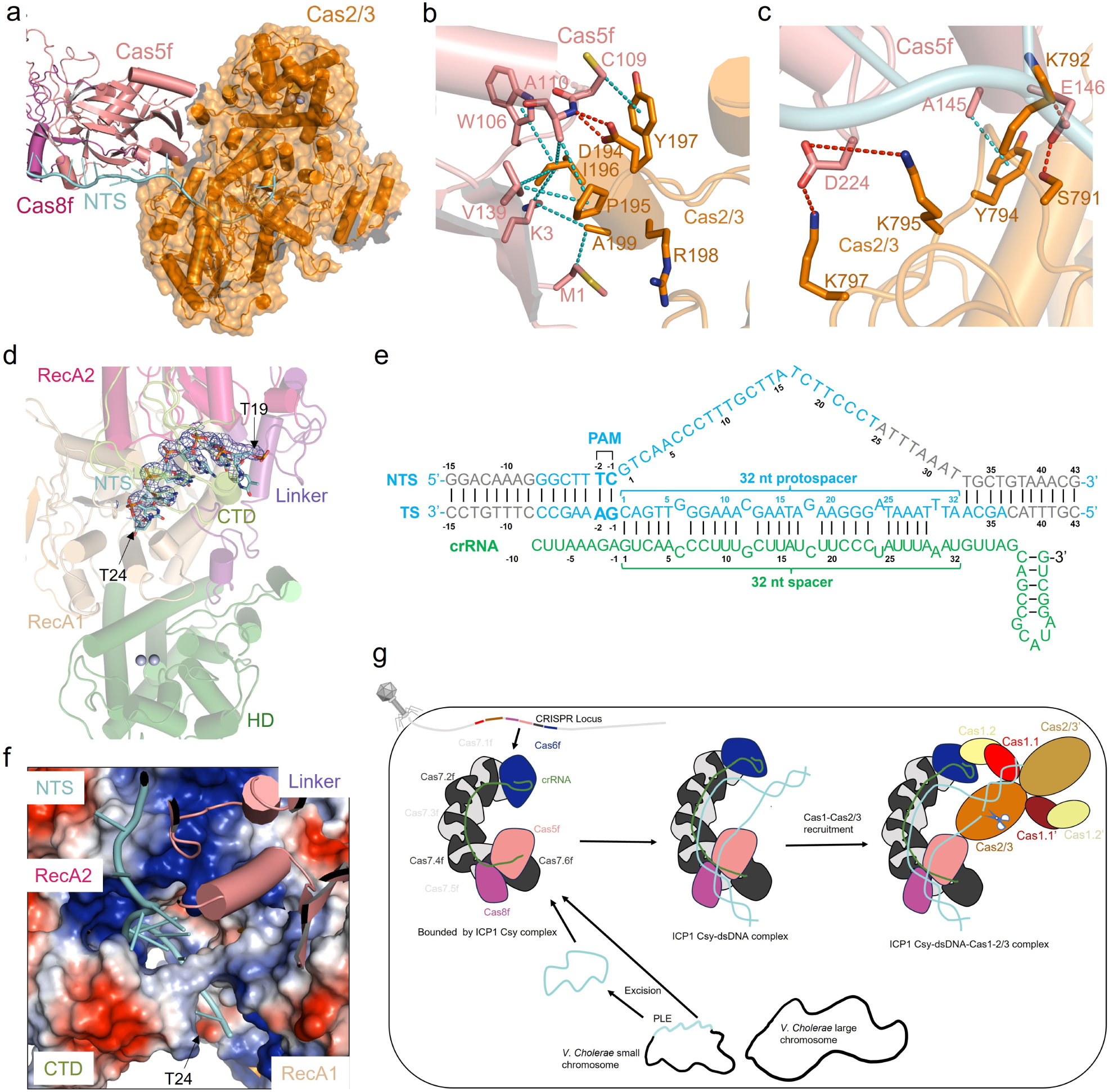
ICP1Cas2/3 binds to Cas5f and captures extended NTS. **a,** ICP1Cas2/3 binds to Cas5f and NTS. The proteins are colored as in Figure 3a. **b,** A short α helix in the HD domain (194-199 residues) of ICP1 Cas2/3 is involved in Cas5f binding. Hydrogen bond and electrostatic interactions are shown as red dashed lines, and hydrophobic interactions are shown as teal dashed lines. The residues in the short α helix primarily bind to residues M1/K3/W106/C109 in Cas5f through hydrophobic interactions. **c,** A loop in the linker domain (791-797 residues) of ICP1 Cas2/3 is involved in Cas5f binding. **d,** Views of the Cas2/3-NTS interaction in the helicase regions. The densities corresponding to T19 to T24 in the NTS are shown in mesh, with protein in cartoon representation. **e,** Schematic drawing of the full R-loop construct with dsDNA and crRNA in the Csy-dsDNA-Cas1-Cas2/3 complex. Nucleotides with no clear density are shown in gray. **f,** Electrostatic surface of the NTS binding channel. The DNA strand is encircled by the positively charged channel formed by the CTD, linker and RecA2 domains of Cas2/3. **g,** Model of the interference mechanism of ICP1 type I-F CRISPR-Cas. Please see text for details.

Through interaction with Cas5f, the Cas2/3 protein is able to capture the separated NTS (Figure 6a). The density allows us to model 31 nucleotides (24 nucleotides in the protospacer region) in the NTS (Figure 6d, e). After the PAM sequence, the NTS traverses the positively charged groove formed by Cas8f (Extended Data Figure 6d), passes over Cas5f and then enters Cas2/3 through a positively charged channel encircled by the CTD, linker and RecA2 domains of Cas2/3 (Figure 6f). Then the NTS crosses the RecA1 domain and extends towards the HD nuclease domain. Consistently, the major cleavage product in a shorter reaction time is ∼40 nt from the 5’ end, suggesting that the complex structure represent a DNA cleaving active state (Figure 1c). Afterwards, the NTS is cleaved progressively, supported by the presence of the bands of lower molecular weight in the gel of *in vitro* cleavage assay (Figure 1c). Consistently, the NTS is bound by the helicase domain of Cas2/3, suggesting subsequent ATP-dependent processive degradation.

## Discussion

Cas1, traditionally known for its role in adaptation ^4^, unexpectedly plays a crucial role in the interference stage of the ICP1 type I-F CRISPR-Cas system, as revealed by biochemical, structural, and *in vivo* assays. Unlike other type I systems, Cas2/3 can only be recruited by the Csy-dsDNA-Cas1 complex. We present 10 structures, including the first snapshots of a Cas1-mediated interference stage and a complete complex for the interference stage of a type I-F system. These findings shed light on the mechanism of interference in the ICP1 CRISPR-Cas system, which lacks the Cas2/3 recruitment domain in the CRISPR Cascade. Interestingly, similar short Cas8 proteins are found in other phages and bacteria(Supplementary Tables 3-4), suggesting a broader presence of Cas1-dependent mechanisms in CRISPR-Cas systems beyond V. cholerae ICP1 and related phages.

Notably, two forms of Csy-dsDNA-Cas1-Cas2/3 complex structures were resolved: a fully assembled symmetrical form and its “half” form. We suggest the fully assembled form may result from high Csy-DNA complex concentrations in incubation samples, not reflecting the transient nature in cells. *In vivo*, we propose the “half” form as predominant. Our interference model for ICP1 type I-F CRISPR-Cas involves initial binding of ICP1 Csy to target dsDNA, followed by recruitment of ICP1 Cas1-Cas2/3 through interactions with Csy-dsDNA. This leads to a conformational change in Cas1-Cas2/3, with NTS extending into Cas2/3’s active site, facilitating target DNA cleavage (Figure 6g).

Zhang et al. recently reported that the ICP1 Csy complex alone has limited binding to dsDNA targets ^36^. However, our MST and ITC assays demonstrate successful binding of ICP1 Csy to dsDNA, albeit with weaker affinity than *Pae* Csy. Additionally, we confirmed this binding through structural analysis (Supplementary Table 2; Extended Data Figure 4). While Zhang et al. suggested that Cas2/3 enhances dsDNA binding and cleavage^36^, we found that ICP1 Csy and Cas2/3 are insufficient for target DNA cleavage without Cas1. This discrepancy may stem from differences in reaction conditions. Our *in vivo* interference and phage assays support the essential role of Cas1 in interference. Furthermore, in our structural analysis, Cas1 bridges the Csy-dsDNA complex to Cas2/3, supporting its role in interference (Figure 6g). Thus, we propose that Cas1 is crucial for the interference stage of ICP1 CRISPR-Cas system *in vivo*, while under high protein concentrations, Cas2/3 and Csy complex alone can cleave target dsDNA in vitro.

CRISPR adaptation occurs in two modes: primed acquisition, where Cas1-Cas2 captures and integrates DNA ends generated during interference; and naive acquisition, where spacers are acquired from unrelated DNA^42^. Primed adaptation, more efficient than naive, involves coupling interference and adaptation processes ^43, 44^. Fusion of Cas2 and Cas3 domains in type I-F systems exemplifies this association, with interference-driven spacer acquisition playing a significant role ^45^. Our study on the ICP1 type I-F system reveals a unique model where Cas1 mediates both interference and adaptation stages, facilitating rapid capture of Cas2/3 products by the Cas1-Cas2/3 complex for efficient primed adaptation. This novel mechanism expands our understanding of CRISPR-Cas systems and offers insights for their application in genome manipulation and beyond.

## Acknowledgements

We thank Prof. Jianlin Lei and Fan Yang (Tsinghua University), Prof. Peiyi Wang and Xiaomin Ma (Southern University of Science and Technology) for EM data collection. We thank the Tsinghua University Branch of the China National Center for Protein Sciences (Beijing) and Southern University of Science and Technology for providing the cryo-EM facility support. We thank the staff at beamlines BL17U1 and BL19U1 of the Shanghai Synchrotron Radiation Facility for their assistance with data collection. We thank the Tsinghua University Branch of China National Center for Protein Sciences Beijing and Dr. Shilong Fan for providing facility support for X-ray diffraction of the crystal samples. We would like to thank Mrs. Wu Yao (State Key Laboratory of Plant Genomics, Institute of Microbiology, Chinese Academy of Sciences) for assistance in the microscale thermophoresis (MST) experiment. This work was supported by the National key research and development program of China (2022YFC2104800, 2022YFC3401500 and 2022YFA1302701), National Natural Science Foundation of China (32371329, 32171274 and 32030056), the Tsinghua-Foshan Innovation Special Fund, No. TFISF-2022THFS6122, the King Abdullah University of Science and Technology (KAUST) Office of Sponsored Research (OSR) under Award, No. OSR-2020-CRG9-4352, Beijing Nova program (20220484160), the Fundamental Research Funds for the Central Universities (QNTD2023-01), China Postdoctoral Science Foundation (2023M740203 and 2023TQ0019) and Postdoctoral Fellowship Program of CPSF (GZB20230051 and GZC20230208).

## Author Contributions

Y. F. conceived and supervised the project. H. W., X. C., Z. G., Z. L. and F. L. purified the proteins, prepared the protein-nucleic acid complexes, performed the activity analysis and binding assays supervised by Y. F. and Y. Z.. L. Z. prepared the cryo-EM samples, collected the cryo-EM data and solved the cryo-EM structures supervised by M. Y.. H. W. collected the diffraction data of crystals. J. Z. solved the crystal structures supervised by J. W.. Y. F. helped solve the crystal structures. Y. F. analyzed the data and wrote the paper with assistance from all the authors.

## Materials and Methods

### Protein Expression and Purification

Recombinant proteins were all overexpressed in *Escherichia coli* BL21 (DE3) strain in Lysogeny broth (LB) medium or M9 for selenomethionine-derivatized proteins. After growth at 37℃, the cells were induced by 0.2 mM isopropyl-β-D-thiogalactopyranoside (IPTG) when the cell density reached an OD_600nm_ of 0.8. The full-length *cas* genes from the *Vibrio cholera* phage ICP1_2011_A were synthesized and inserted into different expression vectors. For the expression and purification of the CRISPR-Cas surveillance complex, *cas8f* and *cas5f* genes were subcloned into pRSFDuet-1 vector (Novagen), in which neither of the two proteins carried a tag. *Cas7f* and *cas6f* genes were subcloned into pETDuet-1 vector (Novagen), in which Cas7f carried His tag. The synthetic CRISPR array was inserted into the second MCS of pACYCDuet-1 vector (Novagen). All the three vectors were co-transformed into *E. coli* BL21(DE3) strain and co-expressed as described above. Cells were harvested by centrifugation, resuspended in lysis buffer (50 mM Tris pH 8.0, 300 mM NaCl, 30 mM imidazole and 5% glycerol), and lysed by sonication. The cell lysate was centrifuged at 20,000 g for 45 min at 4℃ to remove cell debris. The supernatant was loaded to 5 mL HisTrap Fast flow column (GE Healthcare) pre-equilibrated in lysis buffer. The column was washed with 15 column volumes of lysis buffer and 15 column volumes of 95% lysis buffer + 5% eluant buffer (50 mM Tris pH 8.0, 300 mM NaCl, 500 mM imidazole and 5% glycerol), and the intact complex was eluted under an imidazole gradient with its concentration from 53.5 mM to 500 mM. After that, the target protein complex was further subjected to anion exchange chromatography (Buffer-QA: 25 mM Tris pH 8.0, 300 mM NaCl, 2 mM DTT. Buffer-QB: 25 mM Tris pH 8.0, 1 M NaCl, 2 mM DTT). And then, the eluant was finally subjected to two rounds of gel filtration chromatography (Superdex200 10/300 GL, GE Healthcare) in buffer A containing 10 mM Tris pH 8.0, 200 mM NaCl and 5 mM DTT to get rid of the excessive Cas7f protein.

The recombinant Cas7f protein was expressed by the same method as described above. *Cas7f* was subcloned into pETDuet-1-MCS1. The protein was purified by Ni-affinity column chromatography and anion exchange chromatography, and was further subjected to gel filtration chromatography (Superdex200 10/300 GL, GE Healthcare). To obtain the Cas8f-Cas5f complex, *cas8f* and *cas5f* genes were subcloned into pRSFDuet-1 vector (Novagen), in which Cas8f was attached with N-terminal His tag. The protein was purified similarly as the Cas7f protein, but the gel filtration chromatography buffer contained 10 mM Tris pH 8.0, 500 mM NaCl and 5 mM DTT. For the selenomethionine (SeMet) derivatives of Cas8f-Cas5f and the Csy complex, the *E. coli* BL21 (DE3) cells were grown in M9 minimal medium supplemented with 60 mg/L SeMet (Sigma-Aldrich) and specific amino acids: Ile, Leu and Val at 50 mg/L; Lys, Phe and Thr at 100 mg/L. The SeMet protein and mutant proteins were purified as described above.

Template CRISPR sequence: TAATACGACTCACTATAGGGTTAGCAGCCGCATAGGCTGCTTAAAGAGTCAACCCTTTGCTTATCTTCCCTATTTAAATGTTAGCAGCCGCATAGGCTGCTTAAAGAGTCAACCCTTTGCTTATCTTCCCTATTTAAATGTTAGCAGCCGCATAGGCTGCTTAAAGAGTCAACCCTTTGCTTATCTTCCCTATTTAAATGTTAGCAGCCGCATAGGCTGCTTAAAGAGTCAACCCTTTGCTTATCTTCCCTATTTAAATGTTAGCAGCCGCATAGGCTGCTTAAAGAGTCAACCCTTTGCTTATCTTCCCTATTTAAATGTTAGCAGCCGCATAGGCTGCTTAAAGAGTCAACCCTTTG CTTATCTTCCCTATTTAAATGTTAGCAGCCGCATAGGCTGCTTAAAGAGTCAACCCTTTGCTTATCTTCCCTATTTAAATGTTAGCAGCCGCATAGGCTGCTTAAAGA

*Cas1* and *cas2/3* genes were subcloned into pRSFDuet-1-MCS1 (Novagen) and pGEX6p-1 (Novagen), respectively. A TEV protease cleavage site was inserted between the His tag and *cas1* for easy removal of the tag. The lysis buffer for Cas1 contained 1×PBS, 0.5 M NaCl, 5% glycerol, and 30 mM imidazole, and Ni-NTA affinity chromatography was used for purification. The protein was washed with the lysis buffer and digested by TEV Protease in the lysis buffer overnight at 4℃. The eluted Cas1 without tag was concentrated and further purified by gel filtration chromatography (Superdex200 10/300 GL, GE Healthcare) in buffer B containing 10 mM Tris pH 8.0, 0.5 M NaCl and 5% glycerol.

The lysis buffer (buffer C) for Cas2/3 contained 1×PBS, 0.5 M NaCl and 5% glycerol, and GST affinity chromatography was used for purification. The protein was washed with the lysis buffer, and PreScission protease was used to remove the GST tag. The tag-free Cas2/3 was eluted, concentrated, and further purified by gel filtration chromatography (Superdex200 10/300 GL, GE Healthcare) in buffer C.

For expression of the Cas1-Cas2/3 complex*, cas1 and cas2/3* genes were subcloned into pRSFDuet-1 (Novagen), in which Cas1 carried His tag. The lysis buffer for the Cas1-Cas2/3 complex was the same as Cas1, and Ni-NTA affinity chromatography was used for purification. The protein was washed with the lysis buffer and eluted with 1×PBS, 0.5 M NaCl, 5% glycerol and 300 mM imidazole. The eluted Cas1-Cas2/3 complex was concentrated and further purified by gel filtration chromatography (Superdex200 10/300 GL, GE Healthcare) in buffer C. The Cas1-Cas2/3 complex with Cas1 or Cas2/3 mutations were purified similarly as the wild-type complex.

### Crystallization, Data Collection, and Structure Determination

Crystallization of Cas7f, Cas8f-Cas5f, and the Csy complex were performed using the hanging drop vapor diffusion method at 20℃. Crystals of Cas7f was grown from drops consisting of 1 μL protein solution (about 12.5 mg/mL) and 1 μL reservoir solution containing 0.1 M Magnesium chloride hexahydrate, 0.1 M Tris pH 8.5, 33 % v/v PEG 400, 0.1 M Sodium chloride. Crystals of Cas8f-Cas5f was grown from drops consisting of 1 μL protein solution (about 30 mg/mL) and 1 μL reservoir solution containing 0.1 M HEPES pH 7.4∼7.8, 5 % MPD and 8 %∼13 % PEG 6000. Crystals of the Csy complex was grown from drops consisting of 1 μL protein solution (about 20 mg/mL) and 1 μL reservoir solution containing 0.22-0.36 M NH_4_H_2_PO_4_, 0.1 M Sodium acetate trihydrate pH 4.8. The crystals of Cas7f, Cas8f-Cas5f, and the Csy complex were cryo-protected by the reservoir solution supplemented with 10%, 15%, and 35% glycerol, respectively. All the data were collected at SSRF beamlines BL17U1 and BL19U1, integrated and scaled using the HKL2000 package ^46^. The statistics of the diffraction data are summarized in Table S1.

The structure of Cas7f was solved by the molecular replacement (MR) method using PhaserMR in CCP4 with Cas7f in the structure of the Csy-AcrF1-AcrF2 complex (PDB code, 5ZU9) as a search template. The structure of Cas8f-Cas5f was solved by the single-wavelength anomalous dispersion (SAD) method using PHENIX ^47^. The structure of the Csy was solved first by molecular replacement using Cas7f and Cas8f-Cas5f as searching models and the phasing was improved by the data from the crystal of the SeMet protein. Model building was performed using COOT ^48^. The structural models were refined using PHENIX ^47^. The statistics of the structure refinement and the quality of the final structure models are also summarized in Table S1.

### Cryo-EM sample preparation and data acquisition

For the ICP1 Csy-dsDNA-Cas1-Cas2/3 complex, the ICP1 Csy complex and target 58 bp-dsDNA were incubated at a molar ratio of 1:1.5 at 37°C for 2 hours in an incubation buffer of 20 mM HEPES pH 7.5, 150 mM NaCl, and 5% glycerol. ICP1 Cas1 and Cas2/3 were then added at 20-fold excess to the ICP1 Csy complex, and KCl was supplemented to a final concentration of 0.3 M, followed by incubation on ice for 12 hours. Most of the unbound ICP1 Cas1 and Cas2/3 were removed by Ni-NTA affinity chromatography pull-down, and the eluted sample was concentrated to 2 ml. The purified sample was further separated by HiLoad 16/600 Superdex 200 pg (GE Healthcare, 120 mL) (20 mM HEPES pH 7.5, 300 mM KCl) to a well-purified state suitable for cryo-EM of the ICP1 Csy-dsDNA-Cas1-Cas2/3 complex. For the ICP1 Csy-dsDNA (partial duplex) complex, the ICP1 Csy complex and target dsDNA (partial duplex) were incubated at a molar ratio of 1:1.5 om ice for 2 hours, and then was purified by Superdex200 10/300 GL (GE Healthcare, 24 mL) (20 mM HEPES pH 7.5, 150 mM KCl). DNA strands were purchased from Sangon and dissolved in sterile water. Target and non-target DNA strands were mixed together with a molar ratio of 1:1.5, denatured at 95℃ for 5 min, and then annealed by slowly cooling to room temperature.

S8-58 nt target DNA sequence: CGTTTACAGCAATTTAAATAGGGAAGATAAGCAAAGGGTTGACGAAAGCCCTTTGTCC

S8-58 nt non-target DNA sequence: GGACAAAGGGCTTTCGTCAACCCTTTGCTTATCTTCCCTATTTAAATTGCTGTAAACG

S8-49 nt target DNA sequence: ATTTAAATAGGGAAGATAAGCAAAGGGTTGACGAAAGCCCTTTGTCCCT

S8-27 nt non-target DNA sequence: GGCTTTCGTCAACCCTTTGCTTATCAA

Freshly continuous thin layer of carbon grids (Quantifoil 300-mesh Au R1.2/1.3, Micro Tools GmbH, Germany) were glow discharged for 30 s. The grids were then used to apply 4 μL aliquots of samples (3 mg/mL) and blotted for 1.5 s before being plunged into liquid ethane in 100% humidity at 8 °C using a Mark IV Vitrobot (Thermo Fisher Scientific). The datasets of Csy-DNA complex samples were collected using a Titan Krios microscope (Thermo Fisher Scientific) with K2 Summit direct electron detectors and a GIF Quantum energy filter (Gatan). SerialEM software was used to collect data at a nominal magnification of 105,000 × in super-resolution mode with a slit width of 20 eV, and the defocus range was set from −1.5 μm to −2.3 μm. 5,303 movie stacks were obtained with a total dose of 50 e/Å^2^ and an exposure time of 8 s, each micrograph stack contains 32 frames ^49^.

The datasets of ICP1 Csy-dsDNA-Cas1-Cas2/3 complexes samples were collected on a Titan Krios microscope operated at a voltage of 300 kV by a Gatan K3 Summit direct electron detector and a GIF Quantum energy filter (Gatan) with a slit width of 20 eV at a nominal magnification of 81,000 × with a pixel size of 1.1 Å and the defocus range was from −1.3 μm to −1.7 μm. 30,289 movies were collected in three batches using SerialEM software with a total dose of 50 e/Å^2^ and an exposure time of 3 s. The datasets of the Cas1-Cas2 complex samples were collected on a Titan Krios microscope operated at a voltage of 300 kV by a Gatan K3 Summit direct electron detector at a nominal magnification of 105,000 × with a slit width of 20 eV. The pixel size was 0.8374 Å/pixel. The defocus range were set from −1.3 μm to −1.5 μm. The electron exposure on the detector was about 50 e-/Å^2^. 4,375 movies were collected with AutoEMation 2 (written by Jianlin Lei).

### Image processing

In general, all movies were motion-corrected and dose weighted using MotionCor2 for the two datasets ^50^. Gctf was used to determine the contrast transfer function (CTF) parameter and produce the CTF power spectrum^51^. Particles were auto-picked on dose-weighted micrographs using Gautomatch (www.mrclmb.cam.ac.uk/kzhang/Gautomatch/). 2D averages templates for auto-picking were generated by manual-picking and 2D classification.

Next, for the dataset of Csy-dsDNA complex, 1,001,197 particles were extracted from 5120 manually selected micrographs. Three rounds of 2D classification were then performed using cryoSPARC v3^52^, resulting in 563,598 good particles that were exported to RELION 3.1^53^. An initial model was generated using the PDB (6NE0) and two rounds of 3D classification were performed^24^. Subsequently, 240,070 particles were subjected to 3D-refine with a soft mask and per-particle CTF refinement. The resulting map was further improved to a resolution of 3.62 Å by post-processing.

For the dataset of ICP1 Csy-dsDNA-Cas1-Cas2/3 complexes, approximately four million particles were picked from 27,885 selected micrographs by blob-picker in RELION 3.1. After multiple rounds of 2D classification, the final dataset of 1,711,370 particles was subjected to ab-initio and heterogeneous refinement with seven classes using cryoSPARC v3, each with different mask diameters. Among the six classes, those with visible secondary structural elements, such as alpha helices in the 2D class averages, were considered good. One class containing 167,619 particles showed clear Cas1-Cas2 complex features. This class was then subjected to ab-initio and non-uniform refinement after 2D classification, yielding 99,345 good particles, which were further refined to a resolution of 3.36 Å. Two other classes showing clear Csy complex features were subjected to ab-initio, heterogeneous refinement, non-uniform refinement, and local CTF refinement. These classes yielded 90,717 and 168,154 particles, respectively, and the final maps were solved at resolutions of 3.54 Å and 3.15 Å. One of the two classes with larger mask diameters was subjected to imposed C2 symmetry, while the other was displayed with half-symmetry. A total of 235,177 good particles were performed non-uniform refinement and local CTF refinement by cryoSPARC v3, resulting in a final map with a resolution of 2.93 Å. Additionally, the particle subtraction function in RELION was applied to the mask surrounding the Cas5f, Cas3, and Cas6f-Cas1, leading to improved final maps with resolutions of 2.91 Å, 3.14 Å and 3.22 Å, respectively. Furthermore, 132,865 C2-like particles were obtained at resolutions of 3.75 Å for C1 symmetry and 3.72 Å for C2 symmetry.

For the datasets of the Cas1-Cas2 complex, 381,316 selected particles were obtained after two rounds of 2D classification. These particles were performed to ab-initio and heterogeneous refinement with three classes. 101,197 good particles were refined and further improved to 3.60 Å resolution. All reported resolutions are based on the gold-standard FSC=0.143 criteria, and the final FSC curves were corrected for the effect of a soft mask by using high-resolution noise substitution. The final density maps were sharpened by B-factors calculated with the RELION post-processing program. The final maps for model building and figure presentation were performed using DeepEMhancer ^54^. All composite maps were generated in UCSF Chimera using ‘vop maximum’ command and used for model building and refinement ^55^. Further information for all samples is provided in Table S2. Local resolution map was calculated using ResMap ^56^.

### Atomic model building and refinement

Atomic models were predicted and modeled by AlphaFold2 ^57^ and PDB: 6NE0. We docked initial models into final maps by using UCSF chimera and manually adjusted and re-built in COOT ^48^. For cross-validation against overfitting, we randomly displaced the atom positions of the final model by up to a maximum of 0.5 Å, and refined against the final maps using real-space refinement module in PHENIX^58^. The stereochemical quality of each model was assessed using MolProbity ^59^. All the figures were created in PyMOL (www.pymol.org), COOT and UCSF Chimera ^55^ and Chimera X ^60^. Statistics for model refinement and validation are shown in Table S2.

### Protein binding assays

ICP1 Csy complex (1 μM) and dsDNA (1.2 μM) were incubated at 37℃ for 30 min in 20 mM HEPES pH 7.5, 150 mM NaCl, 5% glycerol. Then ICP1 Cas1 (2 μM) was added and incubated at room temperature for 20 min. Finally, ICP1Cas2/3 (4 μM) was added and incubated at 4℃ for 120 min. The mixture was separated using 5% native polyacrylamide gel at 4℃ and visualized by Coomassie blue R250 staining. The sample without dsDNA and the *Pae* group were conducted the same as described above.

### Electrophoretic mobility shift assay

We mixed target with non-target DNA strand at a molar ratio of 1:1.5 to form a dsDNA. FAM was labeled on 3’ end of target DNA strand, synthesized by Sangon, Shanghai. Reactions were performed in a 20 μl buffer system containing 1.6/0.8/0.4/0.2/0.1 μM Csy complex and 0.1 μM dsDNA. All binding reactions were conducted at 37℃ for 30 min in the buffer containing 20 mM HEPES pH 7.5, 100 mM KCl and 5% glycerol. Products of the reaction were separated using 5% native polyacrylamide gels and visualized by fluorescence imaging.

### Plasmid interference assay

ICP1 *cas5f-8f* and CRISPR array were constructed into the pACYCDuet-1 vector. ICP1 *cas1* and ICP1 *cas2/3* were constructed into the pRSFDuet-1 vector. The target DNA sequence (showed below) was constructed into the pUC19 vector. These three plasmids were co-transformed into BL21(DE3) cells. The cells were then spread onto a solid culture medium plate containing Ampicillin (Ap)/Kanamycin (Kan)/Chloramphenicol (Cm) antibiotics. After incubation at 37℃ for more than 10 hours, colonies were picked using a 10 μL pipette and inoculated into 3 sterilized LB culture tubes, each without antibiotics. The OD was allowed to reach 0.6 before adding 0.2 mM IPTG and inducing at 37℃ for 5 hours. After induction, a gradient dilution was performed in 10-fold increments, and 40 μL of each dilution was plated onto solid culture medium plates containing Ap/Kan/Cm and Kan/Cm. After incubation at 37℃ for more than 12 hours, the number of colonies on each plate was counted. The ratio of colonies on the Kan/Cm plate to those on the Ap/Kan/Cm plate represents the activity strength of the ICP1 CRISPR-Cas system, with a higher ratio indicating stronger activity. The experiments of the *Pae* groups were performed similarly.

ICP1 target DNA sequence: ATTTAAATAGGGAAGATAAGCAAAGGGTTGACGA

*Pae* target DNA sequence: CAGGTAGACGCGGACATCAAGCCCGCCGTGAAGG

### Microscale Thermophoresis (MST) measurements

All microscale thermophoresis measurements (MST) were performed on a NanoTemper Monolith NT.115 instrument (NanoTemper Technologies, Munich, Germany). using the Standard Treated Capillaries K022 of the supplier. General settings were applied for all MST experiments as follows: manual temperature control: 25℃, LED power was set at 10% for ICP1 Csy and 20% for *Pae* Csy and MST power was set at high. For MST assay, all the proteins were finally purified in buffer (20 mM HEPES pH 7.5, 150 mM NaCl, 5% glycerol). The Csy complexes were fluorescence-labeled using the Protein Labeling Kit RED-NHS 2nd Generation (NanoTemper Technologies) at 10 μM. 50 nM fluorescence-labeled Csy complexes were added in a 1:1 ratio to a 1:2 dilution series with a final concentration of 2 μM down to 0.061 nM for target dsDNA in buffer (20 mM HEPES pH 7.5, 150 mM NaCl, 5% v/v glycerol) supplemented with 0.05% Tween20. Each experiment was conducted at least 3 times and the similar result was obtained each time. Each protein KD value was obtained with a signal-to-noise ratio higher than 10. Datasets were processed with the MO. Affinity Analysis v2.3 software. The analysis of dose-response curves was carried out with Origin 2022.

### *In vitro* DNA cleavage assay

First, 0.05 μM DNA (6-FAM labeled at the 5’ end of the non-target strand) and 0.8 μM ICP1 Csy complex were incubated at 37℃ for 30 min in the reaction buffer containing 20 mM HEPES pH 7.5, 150 mM KCl, 1 mM TCEP and 5% glycerol. And then 0.1 μM Cas2/3 or 0.05 μM Cas1-2/3 (or mutants) was added into the mixture with 4 mM MnCl_2_ and 1 mM ATP. The reaction was further incubated for 10/30/60 min or 60 min (Figure 5E) and quenched with 1% SDS and heating at 97℃ for 5 min. The products were separated by electrophoresis 14% polyacrylamide gels containing 8 M urea and visualized by fluorescence imaging.

### Plaque assays

The related plasmids or their mutants were transformed into *E. coli* strain BL21(DE3) for the plaque assay, bacteria were grown overnight at 37℃ and then induced for 3 h at 37℃ with 0.2 mM IPTG. The induced bacteria were added to the agar before the plates were poured. 400 μL of the bacterial culture was mixed with 5 mL 0.8% LB-agar (pre-heated to 65℃), poured over the top of a 2.5% LB-agar plate, and let to dry for 30 min at room temperature. 10-fold serial dilutions were performed for T4 phages and 2 μL drops were put on the bacterial layer. After the drops dried, the plates were inverted and incubated for 12 h at 37℃. All experiments include three biological repeats, each including technical replicates in triplicate.

T4 target DNA sequence: ATTCGACAGAACCATCTGCGTACATCATAAATGA;

TGATTGAACCGGAGTATGAATTACTCGTTCTGGA

### Bioinformatics studies of short Cas8 protein

Bacterial Csy1 and Cas8 proteins were searched in the UniProt website (https://www.uniprot.org/), proteins with 100-280 residues were filtered, and CRISPRone ^61^ was used to check if the selected species have a complete CRISPR-Cas system. Among the species identified with a complete CRISPR-Cas system, *Raoultella planticola* strain NCTC12998 was additionally predicted to contain 3 Cas8f proteins with length of less than 100 residues. For phage-encoded short Cas8 proteins, NCBI Virus phage nucleotide database was downloaded and analyzed by CRISPRone ^61^ and seqkit ^62^.

### Data and Materials Availability

The atomic coordinates for the crystal structure of Csy complex (PDB: 8K0K), Cas5f-8f (PDB: 8K0H), Cas7f (PDB: 8K0J), cryo-EM structures of the ICP1 Csy-dsDNA complex (partial duplex) (PDB: 8K27), ICP1 Csy-dsDNA complex (form 1) (PDB: 8K28), ICP1 Csy-dsDNA complex (form 2) (PDB: 8K29), ICP1 Cas1-Cas2/3-dsDNA complex (PDB: 8K25), ICP1 Csy-dsDNA-Cas1-Cas2/3 complex (C1, fully assembled form) (PDB: 8K23), ICP1 Csy-dsDNA-Cas1-Cas2/3 complex (C2, fully assembled form) (PDB: 8K24), ICP1 Csy-dsDNA-Cas1-Cas2/3 complex (half form) (PDB: 8K22), Cas1-Cas2-dsDNA subregion in ICP1 Csy-DNA-Cas1-2/3 complex (PDB: 8K21) and ICP1 Cas1-Cas2/3 complex (PDB: 8K26) have been deposited in the Protein Data Bank (www.rcsb.org).

The cryo-EM density maps reported in this study, ICP1 Csy-dsDNA complex (partial duplex) (EMD-36830), ICP1 Csy-dsDNA complex (form 1) (EMD-36831), ICP1 Csy-dsDNA complex (form 2) (EMD-36835), ICP1 Cas1-Cas2/3-dsDNA complex (EMD-36828), ICP1 Csy-dsDNA-Cas1-Cas2/3 complex (C1, fully assembled form) (EMD-36826), ICP1 Csy-dsDNA-Cas1-Cas2/3 complex (C2, fully assembled form) (EMD-36827), ICP1 Csy-dsDNA-Cas1-Cas2/3 complex (half form) (EMD-36825), Cas1-Cas2-dsDNA subregion in ICP1 Csy-DNA-Cas1-2/3 complex (EMD-36824), Csy1 subregion in ICP1 Csy-DNA-Cas1-2/3 complex (EMD-36832), Cas3 subregion in ICP1 Csy-DNA-Cas1-2/3 complex (EMD-36833), Csy4 subregion in ICP1 Csy-DNA-Cas1-2/3 complex (EMD-36834) and ICP1 Cas1-Cas2/3 complex (EMD-36829) have been deposited in the EM Data Bank.

