## Extended Data Figs for "Cas1 mediates the interference stage in a phage-encoded CRISPR-Cas system"

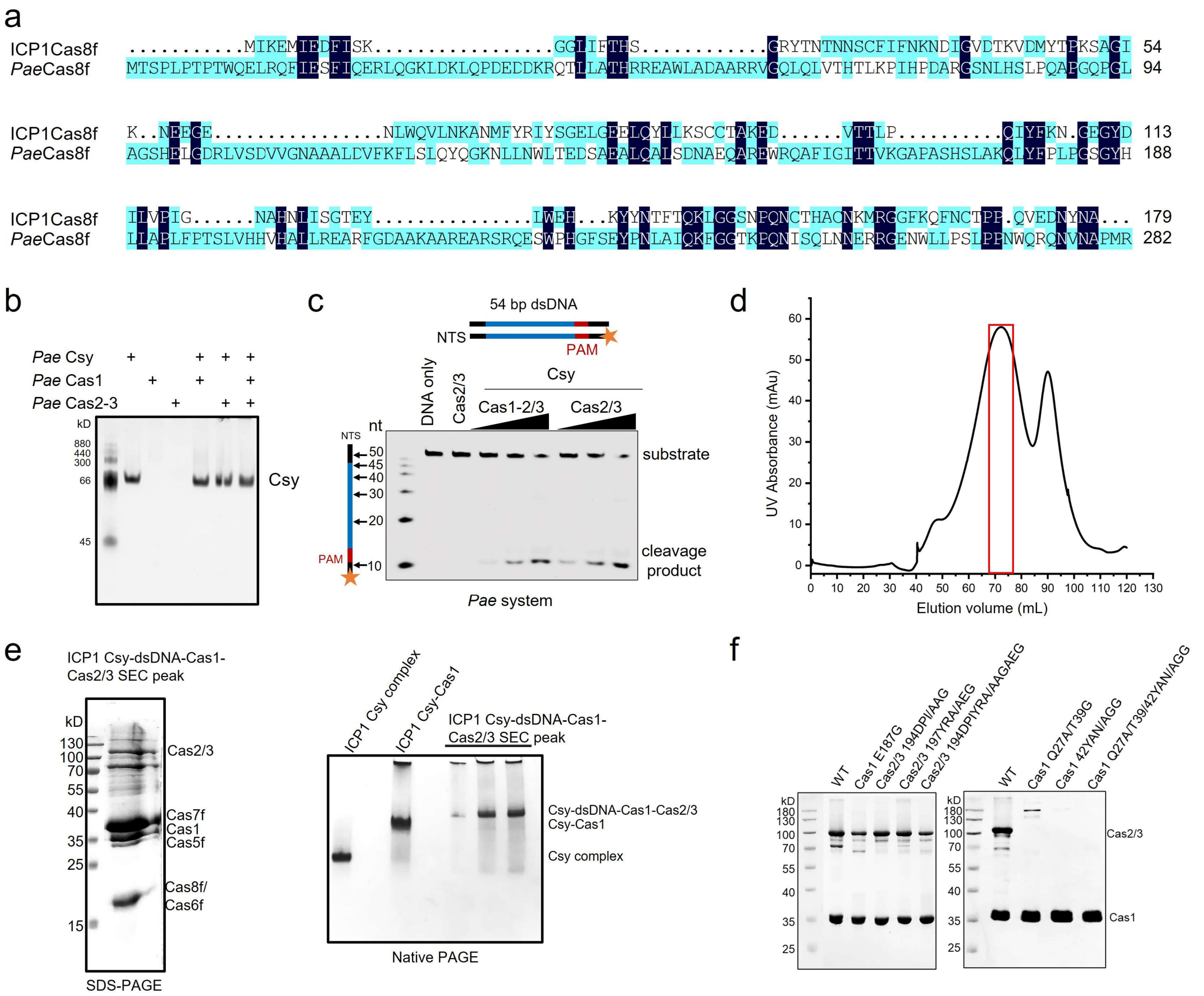

**Extended Data Fig. 1 | ICP1 Cas1 forms a complex with Csy-dsDNA and Cas2/3**

**a,** Sequence alignment of full-length ICP1 Cas8f and *Pae*Cas8f (the region of 1-282 aa). Identical residues are colored in dark blue.

**b,** Protein binding assays. 1 µM *Pae* Csy was incubated with 4 µM *Pae*Cas2/3 only, or first incubated with 2 µM *Pae*Cas1 followed by adding 4 µM Cas2/3. The mixture was separated by native PAGE and visualized by Coomassie blue R250 staining.

**c,** *In vitro* DNA cleavage of the *Pae* CRISPR-Cas system. 0.04 µM dsDNA was preincubated with 0.4 µM Csy complex. Next, Cas1-Cas2/3 (0.04/0.08/0.16 µM) or Cas2/3 (0.08/0.16/0.32 µM) with 1 mM ATP and 5 mM $MgCl_2$, 5 mM $CaCl_2$ and 75 µM $NiSO_4$ were added into the reaction system. The reaction was terminated at 60 min. The products were separated by Urea-PAGE and visualized by fluorescence imaging.

**d,** Gel filtration chromatography of ICP1 Csy-dsDNA-Cas1-Cas2/3 complex.

**e,** Fractions (red box in d) were analyzed by SDS-PAGE and native PAGE in the right panels.

**f,** Purified ICP1 Cas1-Cas2/3 complex and mutants were analyzed by SDS-PAGE.

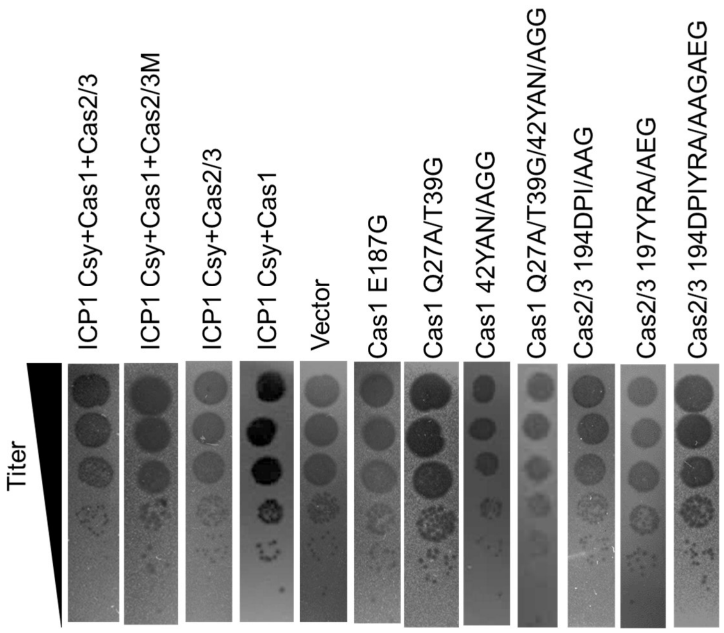


**Extended Data Fig. 2 | Plaque assays of the ICP1 CRISPR-Cas system. Related to Figure 1h and 5d**

The experiment has been repeated independently for 3 times and a representative result is shown. Cas2/3M in this figure represents Cas2/3 D112A/D280A.

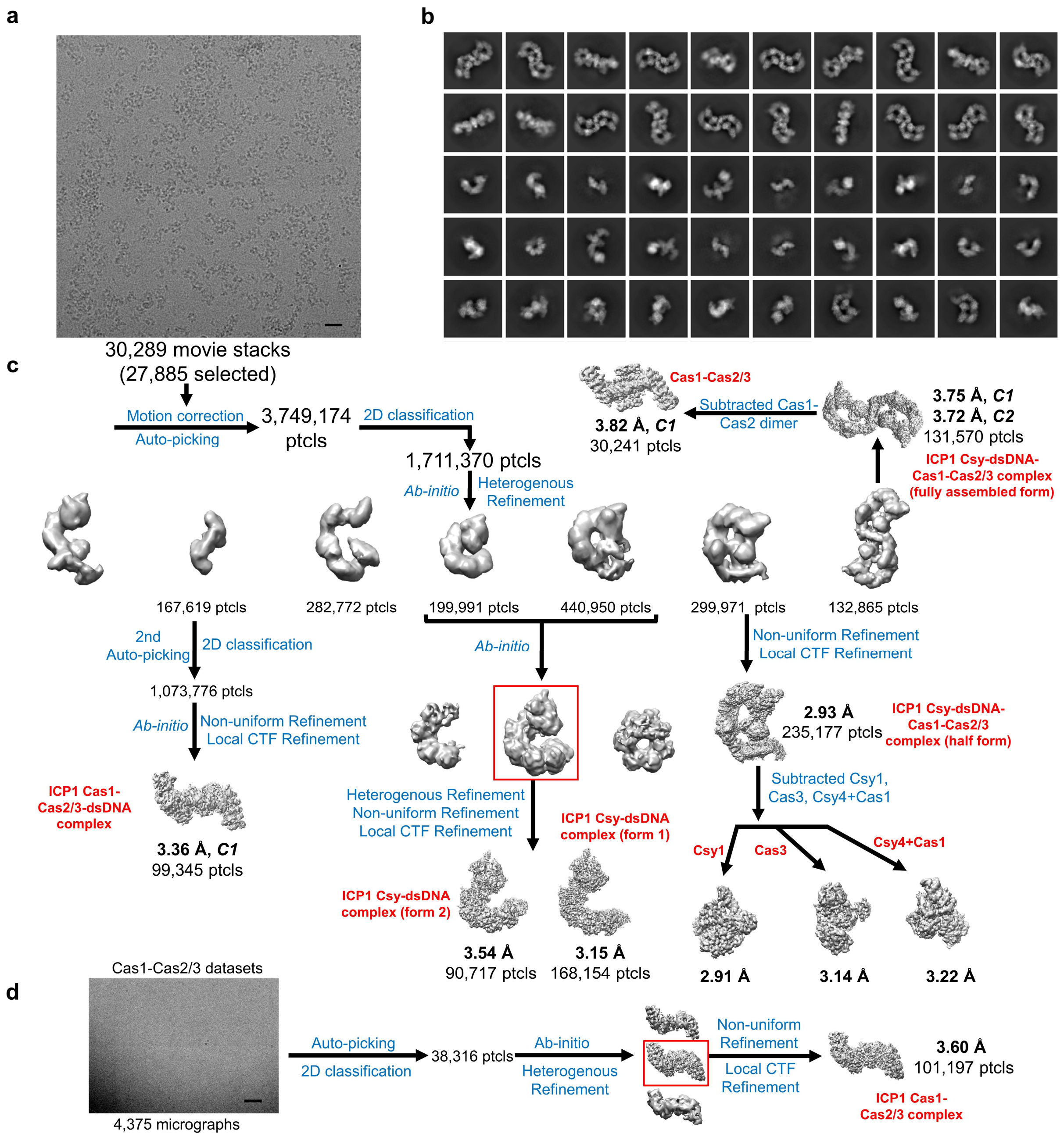


**Extended Data Fig. 3 | Cryo-EM micrograph, 2D class averages and data processing of the ICP1 Csy-dsDNA-Cas1-Cas2/3 complex**

**a,** Representative micrograph of the sample of ICP1 Csy-dsDNA-Cas1-Cas2/3 complex. The scale bar represents 20 nm.

**b,** Representative 2D class averages of the sample of ICP1 Csy-dsDNA-Cas1-Cas2/3 complexes.

**c,** Cryo-EM image-processing flow-chart of ICP1 Csy-dsDNA-Cas1-Cas2/3 complex.

**d,** Image-processing procedure of ICP1 Cas1-Cas2/3 complex. The scale bar represents 10 nm.

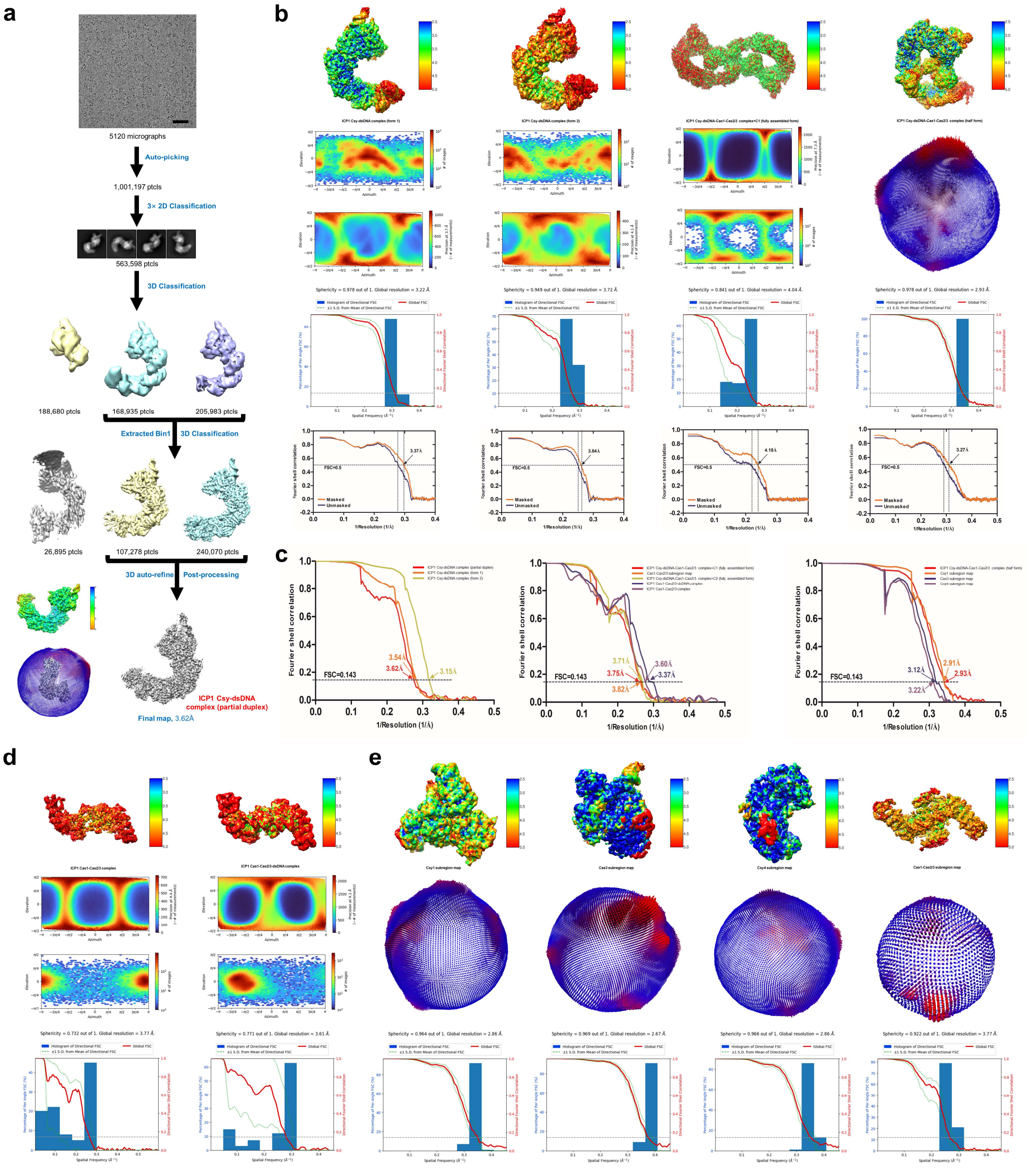


**Extended Data Fig. 4 | Local resolution estimation of the cryo-EM structures of the ICP1 Csy complexes, Cas1-Cas2/3 complexes and Csy-dsDNA-Cas1-Cas2/3 complexes.**

**a,** Overview of data collection and image-processing procedure of ICP1 Csy-dsDNA complex (partial duplex) (see Methods). The scale bar represents 10nm.

**b,** Image processing procedure, local resolution maps, angular distribution, 3DFSC analyses and model-to-map FSC curves (FSC=0.5) were shown for ICP1 Csy-dsDNA complexes (form 1 and form2) and Csy-dsDNA Cas1-Cas2/3 complexes (fully assembled form and half form).

**c,** Gold-standard FSC curves (FSC=0.143) of the final 3D reconstructions of ICP1 Csy complexes, Cas1-Cas2/3 complexes and Csy-dsDNA-Cas1-Cas2/3 complexes.

**d,** Local resolution maps, angular distribution and 3DFSC analyses were shown for the ICP1 Cas1-Cas2/3 and Cas1-Cas2/3-dsDNA complex.

**e,** Local resolution maps, angular distribution and 3DFSC analyses were shown for the subregion maps of Csy1,

Cas3, Csy4 and Cas1-Cas2/3 for ICP1 Csy-dsDNA-Cas1-Cas2/3 complex.

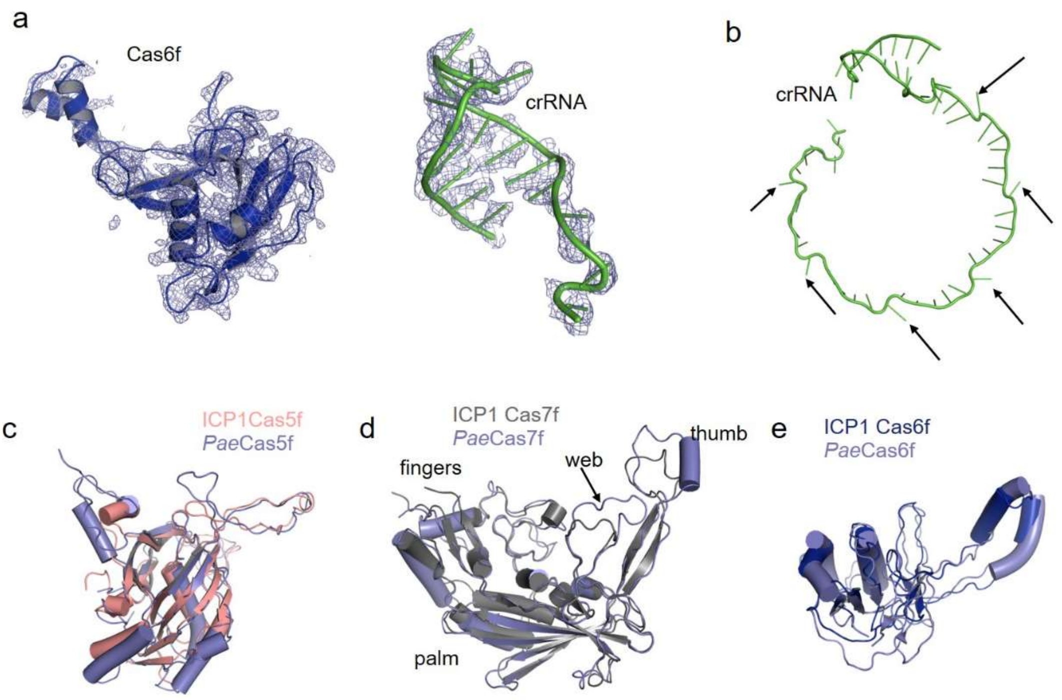


**Extended Data Fig. 5 | Structural characteristics of ICP1 Csy and comparison with *Pae* Csy**

**a,** 2Fo-Fc electron density of the Cas6f subunit and crRNA 3' hairpin in the crystal structure of Csy complex contoured at 1 σ.

**b,** The crRNA in ICP1 Csy exhibits distortions (kinks) along the Cas7f backbone at regular 6-nucleotide intervals.

**c-e,** Structural alignment of ICP1 Cas5f and *Pae*Cas5f (c), ICP1 Cas7f and *Pae*Cas7f (d) and ICP1 Cas6f and *Pae*Cas6f (e). ICP1 Cas7f has shorter "thumbs" and "webs" regions than *Pae*Cas7f (d).

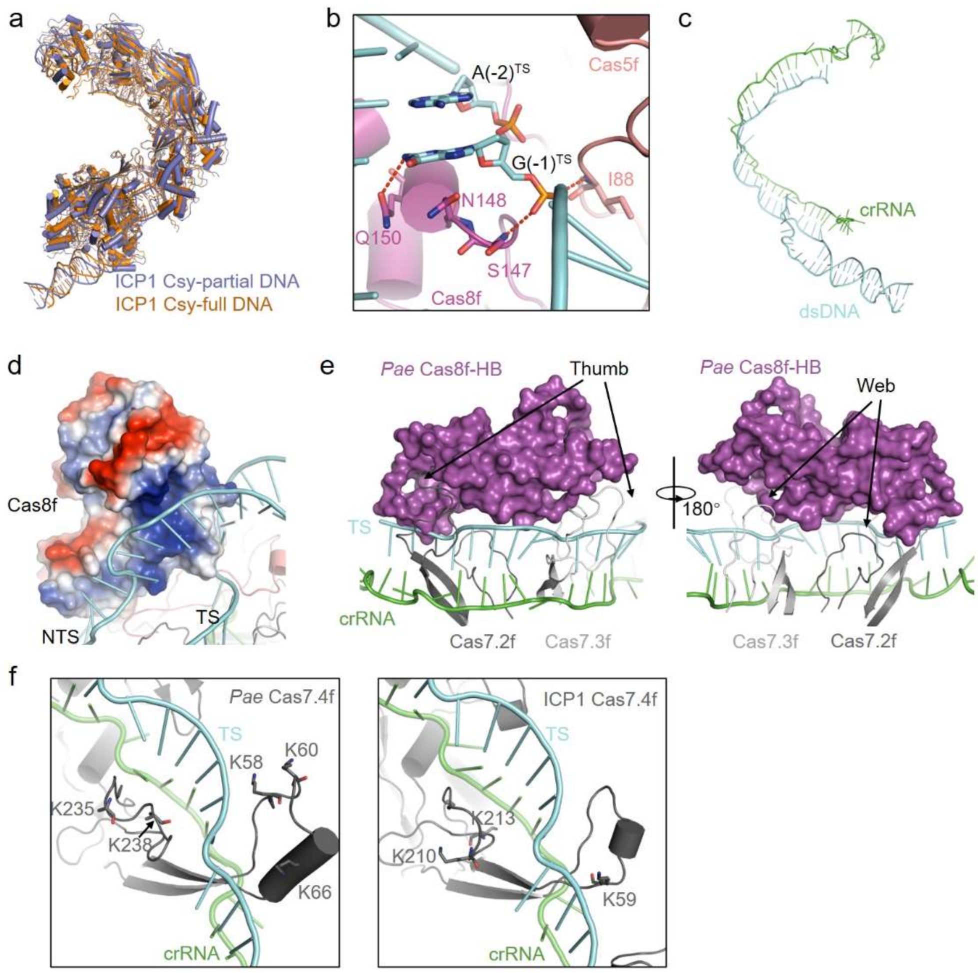


**Extended Data Fig. 6 | Structural characteristics of ICP1 Csy-dsDNA and comparisons with *Pae* Csy-dsDNA**

**a,** Structural alignment of ICP1 Csy-partial dsDNA and Csy-full dsDNA (full R-loop form), they are colored in slate and tv_orange, respectively.

**b,** Details of target DNA PAM recognition in the Cas8f and Cas5f region.

**c,** crRNA:target DNA heteroduplex and duplex DNA are shown in cartoon model. There is one kinked-off nucleotide at every $6^{th}$ position in the crRNA:target DNA heteroduplex.

**d,** NTS is attached to the positively charged groove of the Cas8f subunit.

**e,** Target DNA strand is locked by Cas8f-HB, the “thumb” and “web” regions of Cas7.2f and Cas7.3f in the *Pae* Csy-dsDNA complex (PDB code: 6NE0). Two views are shown.

**f,** K58 and K60 of Cas7f, two critical residues for dsDNA binding in *Pae*Cas7f, is lacking in ICP1 Cas7f.

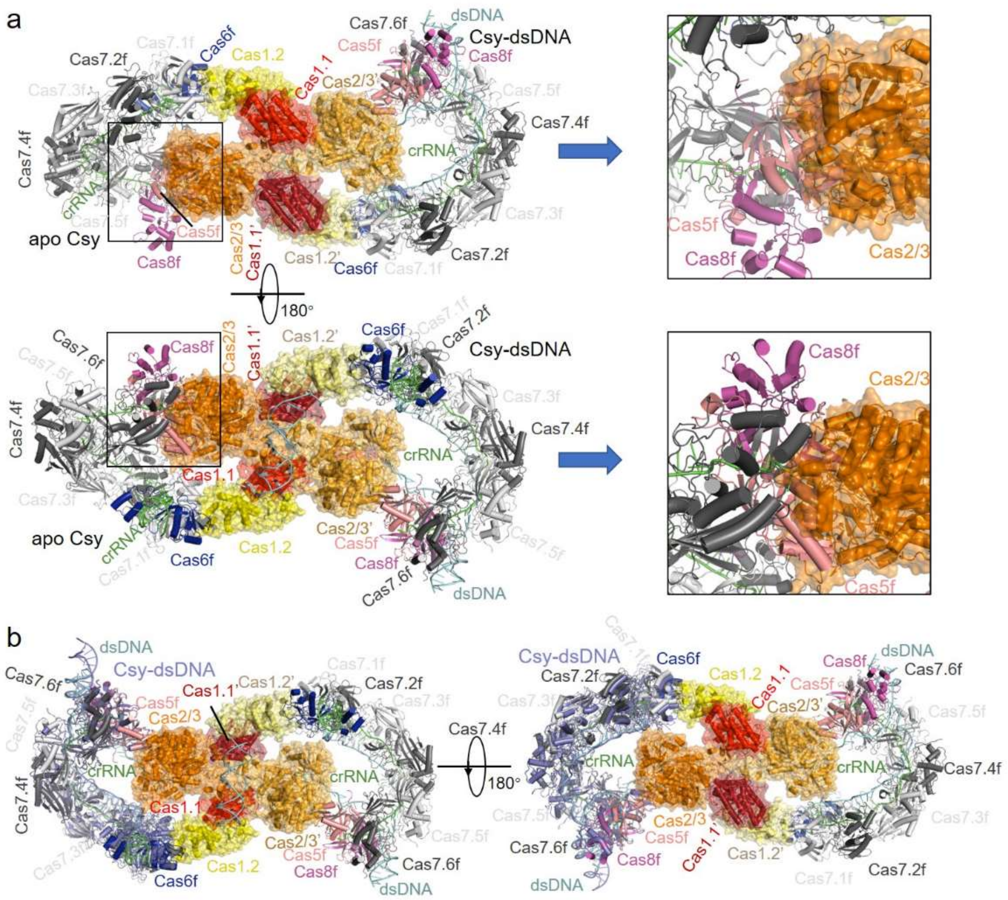


**Extended Data Fig. 7 | Cas1-Cas2/3 binds the Csy-dsDNA form but not the apo Csy**

**a,** Superimposition of the apo Csy onto the Csy within the Csy-dsDNA-Cas1-Cas2/3 complex at the Cas6f subunit (in the left side of the complex). Two views are shown. Close view of the regions in boxes in the left side are shown in the right side.

**b,** Structural alignment between the Csy-dsDNA complex (colored slate) and that within the Csy-dsDNA-Cas1-Cas2/3 complex. Two views are shown.

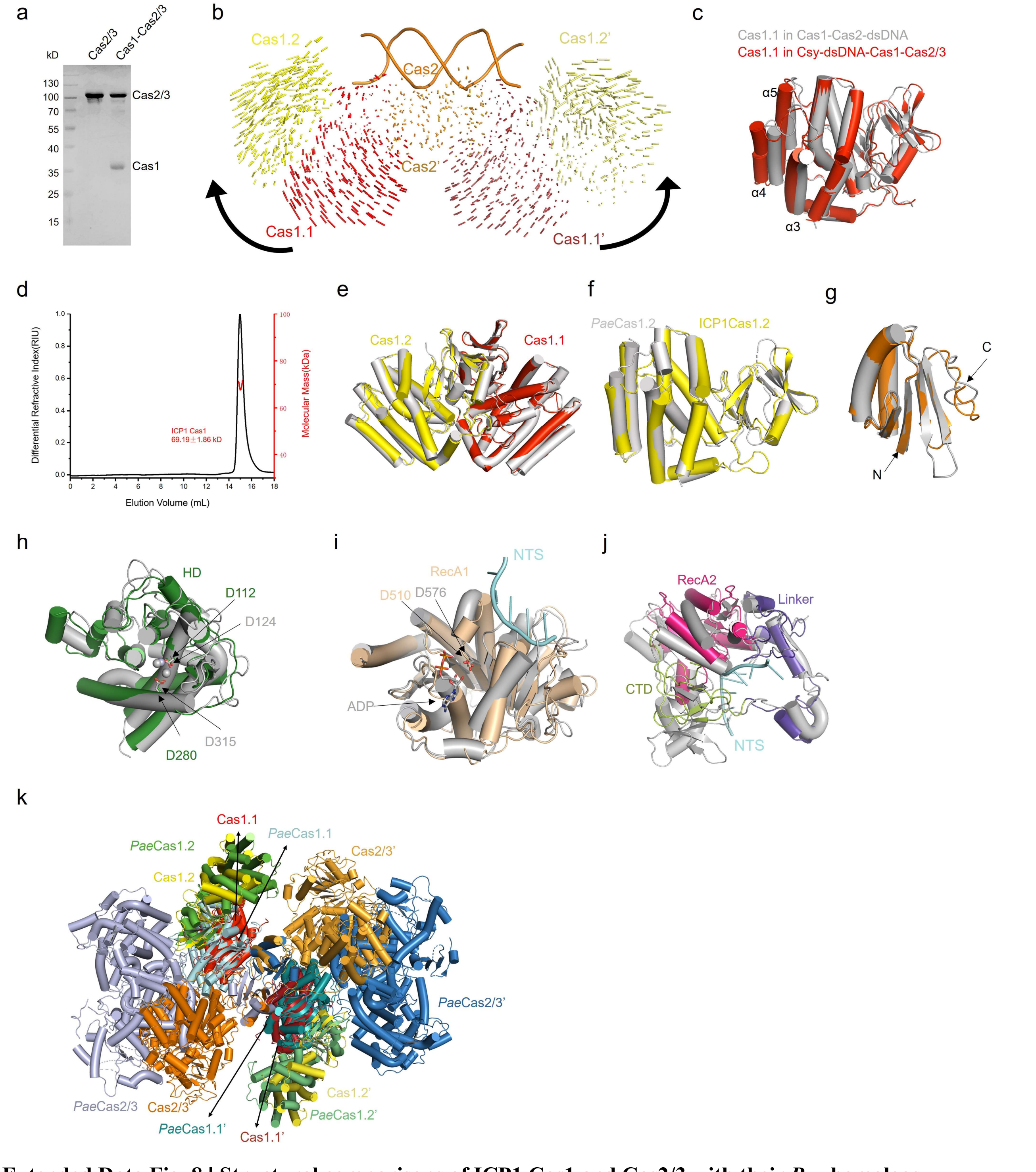


**Extended Data Fig. 8 | Structural comparisons of ICP1 Cas1 and Cas2/3 with their *Pae* homologs**

**a,** Purified ICP1 Cas2/3 and Cas1-Cas2/3 complex were analyzed by SDS-PAGE.

**b,** Structural comparison between apo ICP1 Cas1-Cas2 and that in Cas1-Cas2-dsDNA complex. The Cas2 dimer is superimposed. Vector length correlates with the domain motion scale. The black arrows indicate domain movements within Cas1-Cas2 complex upon dsDNA binding.

**c,** Structural comparison of ICP1 Cas1.1 in Cas1-Cas2-dsDNA complex and that in Csy-dsDNA-Cas1-Cas2/3 complex.

**d,** Static light scattering (SLS) study of ICP1 Cas1. The calculated molecular weight of the main peak is shown.

**e,** Structural comparison of ICP1 Cas1 dimer in Csy-dsDNA-Cas1-Cas2/3 complex and apo Cas1 dimer (PDB code: 4W8K, gray).

**f,** Structural comparison of ICP1 Cas1.2 in Csy-dsDNA-Cas1-Cas2/3 complex and *Pae*Cas1.2 (PDB code: 3GOD, gray).

**g-j,** Structural comparison of the Cas2 domains (g), HD domains (h), RecA1 domains (i), and the region of RecA2/linker/CTD (j) of ICP1 Cas2/3 and *Pae*Cas2/3 (PDB code: 5B7I, gray).

**k,** Structural comparison of ICP1 Cas1-2/3 and *Pae*Cas1-2/3 (PDB code: 8FLJ). Cas2 domain of ICP1 and *Pae* are aligned. The distance between the ICP1 Cas1.1/1.2 and PaeCas1.1/1.2 mass centers is 1.74 Å, and between the ICP1 Cas1.1'/1.2' and PaeCas1.1'/1.2' is 3.78 Å. The distance between ICP1 Cas2/3 and PaeCas2/3 mass centers is 35.75 Å, and between the ICP1 Cas2/3' and PaeCas2/3' is 28.84 Å.

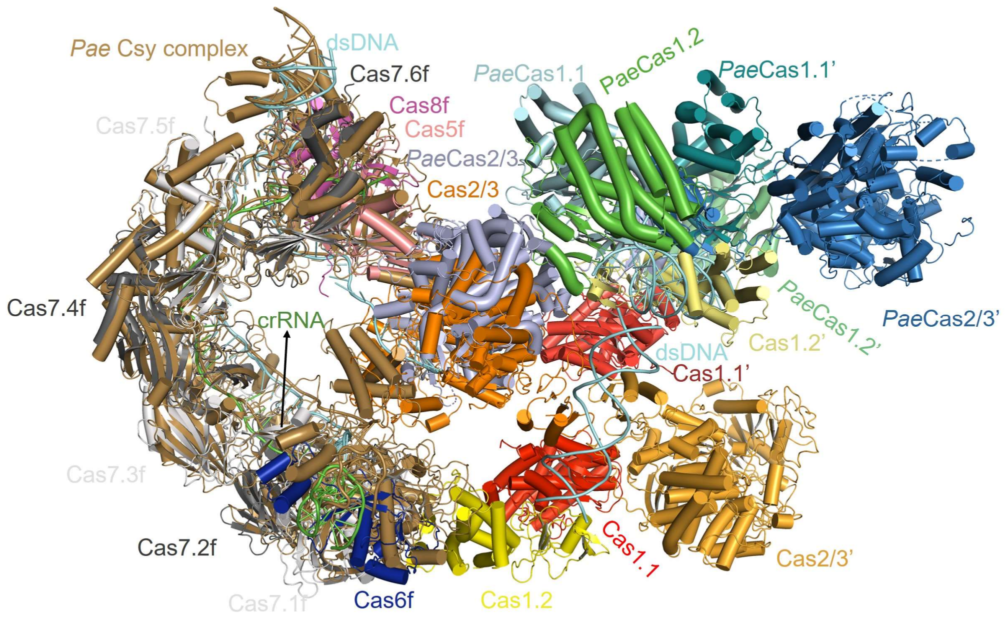


**Extended Data Fig. 9 | Comparison of structure of ICP1 dsDNA-Csy-Cas1-2/3 and the model of *Pae* dsDNA-Csy-Cas1-2/3.**

Superimposition of ICP1 Csy complex and *Pae* Csy complex (PDB code: 6NE0). *Pae*Cas1-2/3 (PDB code: 8FLJ) is docked to the *Pae*Cas8f HB domain.

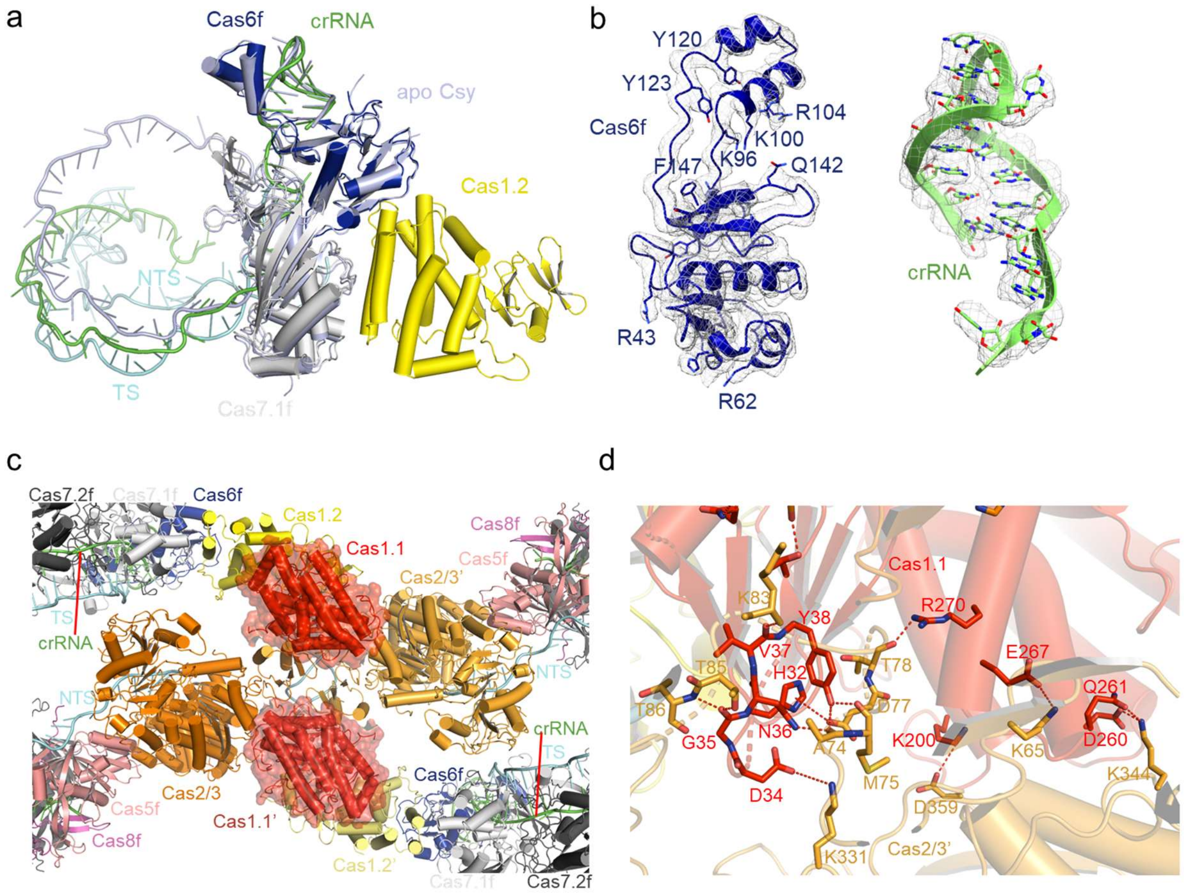


**Extended Data Fig. 10 | Structural details of the Csy-dsDNA-Cas1-Cas2/3 complex**

**a,** Superimposition of the apo Csy onto the Csy within the Csy-dsDNA-Cas1-Cas2/3 complex at the Cas6f subunit. The apo Csy structure is colored in light blue.

**b,** The densities corresponding to Cas6f and crRNA 3' hairpin in the Csy-dsDNA-Cas1-Cas2/3 complex are shown in mesh. Cas6f and crRNA are shown in cartoon and representative residues are shown as sticks.

**c,** Each ICP1 Cas1.1 interacts with both two Cas2/3 protomers in the Csy-dsDNA-Cas1-Cas2/3 complex.

**d,** Detailed interactions between Cas1.1 and Cas2/3'. Hydrogen bond and electrostatic interactions are showed as red dashed lines.
